# Functional Trajectories during innate spinal cord repair

**DOI:** 10.1101/2023.01.31.526502

**Authors:** Nicholas O. Jensen, Brooke Burris, Lili Zhou, Hunter Yamada, Catrina Reyes, Mayssa H. Mokalled

## Abstract

Adult zebrafish are capable of anatomical and functional recovery following severe spinal cord injury. Axon growth, glial bridging and adult neurogenesis are hallmarks of cellular regeneration during spinal cord repair. However, the correlation between these cellular regenerative processes and functional recovery remains to be elucidated. Whereas the majority of established functional regeneration metrics measure swim capacity, we hypothesize that gait quality is more directly related to neurological health. Here, we performed a longitudinal swim tracking study for sixty individual zebrafish spanning eight weeks of spinal cord regeneration. Multiple swim parameters as well as axonal and glial bridging were integrated. We established rostral compensation as a new gait quality metric that highly correlates with functional recovery. Tensor component analysis of longitudinal data supports a correspondence between functional recovery trajectories and neurological outcomes. Moreover, our studies predicted and validated that a subset of functional regeneration parameters measured 1 to 2 weeks post-injury is sufficient to predict the regenerative outcomes of individual animals at 8 weeks post-injury. Our findings established new functional regeneration parameters and generated a comprehensive correlative database between various functional and cellular regeneration outputs.

## Introduction

As vertebrates with elevated regenerative potential, adult zebrafish are an excellent model for innate spinal cord (SC) repair. After complete SC transection, pro-regenerative responses involving the immune system, neurons, and glia contribute to functional regeneration (1). Prompted by a host of injury-induced regeneration promoting factors, glial and axonal bridges reconnect the lesioned tissue, neurons regenerate proximal to the lesion, and swim capacity is restored, all within 6–8 weeks post-injury (wpi) (2). Due to the remarkable ability of adult zebrafish to reverse paralysis, recovery of swim function is an ultimate metric and a central readout of SC repair.

Functional measurements of swim ability may be classified into two broad categories: swim capacity, which represents the amount or extent of swimming a fish can perform under specific conditions; and gait quality, which reflects the quality of a swim bout and the healthiness of a fish during active swimming. One of the earliest assays, in 1997, to evaluate swim function in zebrafish was a binary, recovered or not recovered, quality scoring system assigned by an observer using a startle assay (3). In this assay, fish that respond to a stimulus and swim mostly using their tail instead of their head are labeled as recovered. Based on these and other criteria, swim quality scoring systems on 1 to 5 and 1 to 6 scales have also been developed (4, 5). Two capacity measurements were subsequently introduced: time to exhaustion in 1998 (6) and swim distance in a static tank in 2004 (7). These offered several advantages: they are less subjective, provide continuous scale measurements, and are scalable for longer assays using computational tools. Consequently, swim distance or time to exhaustion have been widely applied to SC regeneration studies between 1997 and 2020. Since 2020, the toolkit to quantify swim function has rapidly expanded and diversified. Traditional and recently established measurements of swim capacity include swim endurance as quantified by time to exhaustion against increasing water current velocities (2, 6, 8, 9, 10), maximum speed either against increasing water current velocities or in a static tank (11, 12, 13, 14), swim distance (7, 9), time active (9, 10), time swimming against flow (9), burst frequency (9, 10), and average position of the fish against a water flow axis (“mean y”) (9, 10). Although new quality measurements have been developed for larval fish including body angle (15), power spectral densities (16), and centerline posture kinematics (17), published quality measurements for adult fish are limited to manual scoring of perceived swim ability (3, 4, 5), tail beat frequency and amplitude (12). Moreover, manually assigned scores are subjective and do not scale well for higher throughput. Thus, behavioral tests to quantify gait quality in adult zebrafish are less developed, yet much needed for accurate assessment of musculoskeletal function in general and of neural regeneration in particular.

Gait analysis offers a scalable approach to quantify swim quality. Prior applications of gait analysis, such as dorsal centerline posture and tail beat kinematics, supported the model that functional recovery emerges directly from structural recovery among ray-finned fishes (12, 17, 18). These studies revealed stepwise recovery of specific gait features in correlation with axon regeneration from V2a interneurons (12, 19). These findings suggested that formerly paralyzed zebrafish may regain the ability to swim in a preinjury-like manner after neural regeneration. Yet, further analysis of swim gait metrics, including posture and behavior, is required to confirm this hypothesis.

A first step in gait analysis is the annotation of elemental components of gait in swim videos: posture and behavior. Automated pose and behavior annotation tools for model organisms are increasingly accessible. While traditional computational methods for zebrafish pose annotation include thresholding and skeletonization (20, 21) or template-fitting (22, 23), many newer cross-species methods prioritize deep learning. Such tools include DeepLabCut (24), DeepPoseKit (25), OptiFlex (26), and SLEAP (27). For zebrafish, quantifying the dorsal centerline is an appropriate representation of body posture. Despite its limitations in measuring vertical position, body deformity in the dorso-ventral axis, or pectoral fin motion, dorsal centerline posture and gait has been applied to larval and adult locomotion analysis (12, 20, 21) and lateral tail beating is the source of most swim propulsion (28). Additionally, certain injury-induced gait alterations manifest in the centerline (12). Similar to recent pose annotation methods, automated behavior annotation is often performed through machine learning either by clustering in a latent space (20, 21, 29), such as the pre-packaged tool B-SOiD (30), or through deep learning, including DeepBhvTracking (31) or DeepEthogram (32). Despite advances in pose and behavior annotation, methods to compare episodes of the same zebrafish behavior, either between animals or between recovery timepoints, are yet to be applied. Comparing multiple episodes of the same behavior is a critical component of gait quality analysis. One viable option includes novelty detection, which classifies behavior episodes as inliers or outliers relative to a control distribution. Another option is posture parameter quantification, which has been used in larval zebrafish (17). Examples of posture parameter quantification from human and mouse studies used behavior and gait features, such as quantified response to stimulus and range or speed of motion, to measure functional recovery (33, 34, 35, 36). As swim quality is proposed to be a more direct readout of SC function and musculoskeletal wellness than swim capacity, recovery metrics based on posture and gait have the potential to become standards for functional assessment.

Given the recent diversification of swim capacity and gait quality measurements, this study compares various established and novel functional recovery metrics and describes their correlations with anatomical regeneration. To this end, we tracked swim function prior to injury and weekly between 1 and 8 wpi. Fish identities were tracked throughout the experiment, and cellular regeneration parameters were measured at 8 wpi. Rostral compensation as established as a new gait feature that highly correlates with functional recovery. For comprehensive analysis of gait features, we also describe recovery trajectories using tensor decomposition of longitudinal recovery data. By observing correlations between established swim capacity and new gait analysis metrics with cellular regeneration features, we propose and validate a strategy for predicting 8 wpi recovery outcomes using 1 and 2 wpi swim function parameters. This study establishes a new gait quality measurement of neuromuscular health, provides a comprehensive correlative analysis of various functional and cellular regeneration outputs, and will help reduce measurement redundancy in future studies.

## Results

### A longitudinal study of spinal cord regeneration parameters

We devised a longitudinal study that allowed us to track cellular and swim recovery metrics in individual animals for 8 weeks of SC regeneration (Figure 1A). Sixty wild-type fish of the Ekkwill/AB strains (30 males and 30 females) were housed separately for the duration of the study. Swim behavior was recorded for each fish prior to injury and weekly between 1 and 8 wpi. At each time point, fish were placed in an enclosed swim tunnel and a dorsally placed camera was used to capture swim behavior at 70 frames/s for a total of 15 min at water current velocities of 0 cm/s (5 min), 10 cm/s (5 min), then 20 cm/s (5 min). At 8 wpi and following swim behavior recording, axon tracing experiments and glial bridging measurements were performed to assess cellular regeneration metrics.

**Figure 1.**
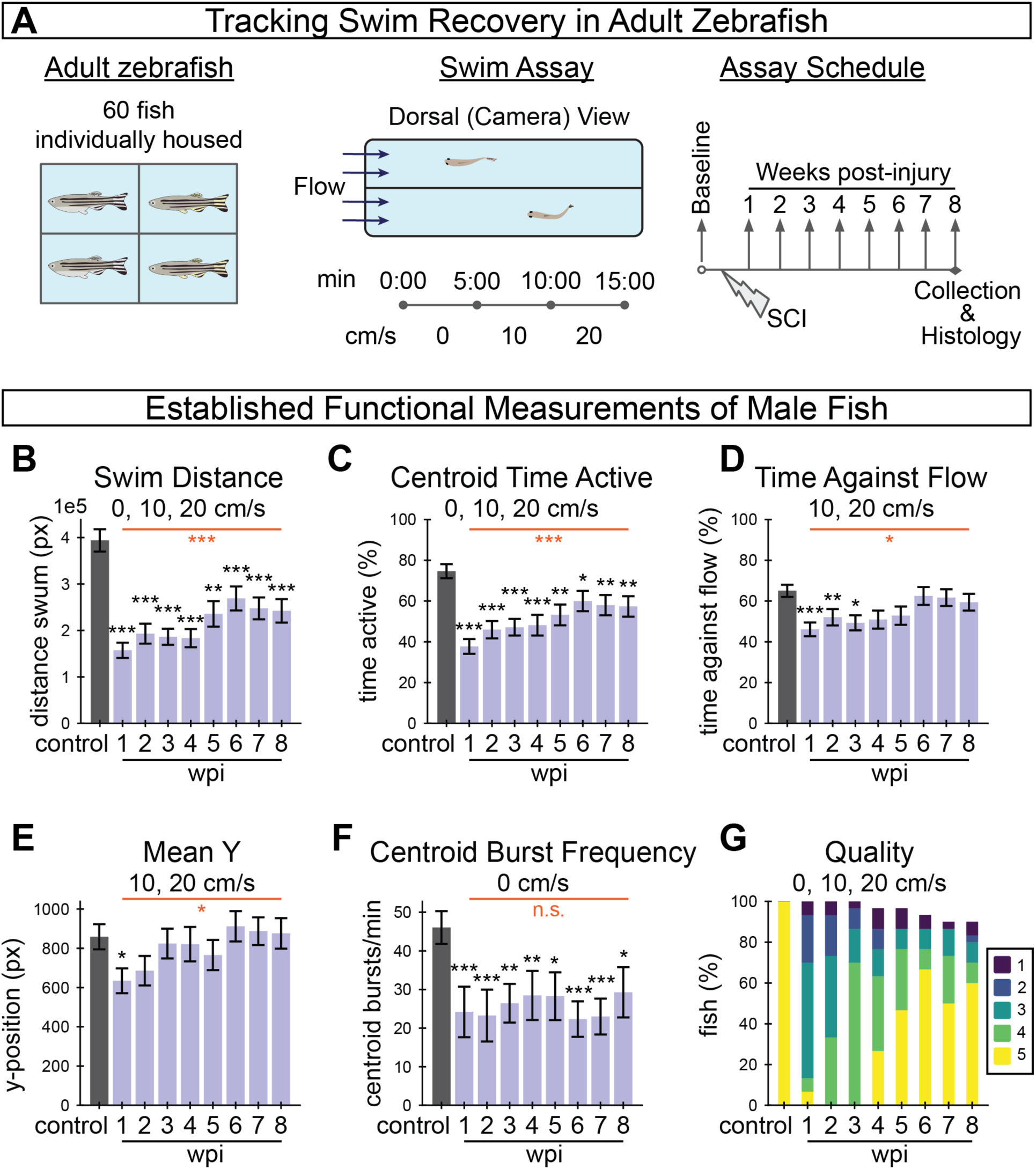
Longitudinal tracking of swim recovery after spinal cord injury in adult zebrafish. **(A)** Schematic representing an overview of the tracking experiment. A total of 60 fish (30 males and 30 females) were housed separately throughout the entire experiment to allow for longitudinal tracking of individual fish throughout the study. Swim behavior was recorded prior to injury, and weekly between 1 and 8 wpi. At each time point, fish were video recorded in a swim tunnel for fifteen minutes. In each 15-min assay, fish were first acclimated in the absence of water current for 5 minutes (0 cm/s), then water current velocity was increased to 10 cm/s at 5 min and to 20 cm/s at 10 min. **(B-G)** A suite of established functional measurements were used to assess swim recovery for male fish following SC transection. The swim recovery outcomes for female fish are shown in Figure S1. Swim distance (B), activity as measured by centroid movement (C), time swimming against the flow (D), mean y-position in the axis of flow (E), and burst frequency as measured by centroid movement (F) represent swim capacity measurements. Perceived swim quality scores are shown in G. Error bars depict SEM and statistical significance was determined by one-way repeated mixed measures ANOVA followed by post-hoc Student’s t-tests. p-value markers in black represent comparisons between each time point post-injury relative to control measurements prior to injury. Red horizontal bars and p-values marked in red show significance between 1 wpi and 8 wpi. ***P<0.001; **P<0.01; *P<0.05; ns, P>0.05.

To begin to assess SC regeneration parameters in this study, we first calculated previously established swim behavior measurements from the captured swim videos (Figure 1B-G). Functional measurements indicated that zebrafish are still capable of recovery from SC transection while isolated from other fish. Swim distance, centroid time active, time against flow and mean y-position decreased with acute injury at 1 wpi relative to controls, then significantly increased between 1 and 8 wpi to approach uninjured control levels (Figure 1B-E). Centroid burst frequency for male fish decreased at 1 wpi but did not show significant improvement at 8 wpi (Figure 1F). Similarly, the proportion of fish that swam in a healthy manner, according to perceptually assessed swim quality, decreased after injury and increased through recovery (Figure 1G). We noted that male fish exhibited this stereotypical recovery trend more strongly than females in this experiment (Figure S1A-F). Overall, the female fish in this experiment regenerated poorly (Figure S1F) and were less energetic both before and after injury, which is not necessarily typical of zebrafish (14). For this reason, female fish data are presented in the supplement.

### Spinal cord injury disrupts healthy swim gait

While established measurements of swim behavior captured the recovery of swim capacity after SCI, our swim videos showed even more pronounced swim gait differences. Comparing one period of oscillatory forward swim behavior (cruise) before injury and at 1 wpi, we found that acutely injured fish swim using more motion in the rostral portion of their body (Figure 2A-B). We thus postulated that a scalable quantification of gait healthiness would provide a markedly improved functional regeneration parameter compared to established swim capacity metrics and to manually assigned quality scores. To perform high-throughput gait analysis, we began by annotating dorsal centerline posture (Figure 2C). After training DeepLabCut (24, 37) to annotate keypoints on the centerline, we filtered and processed the poses, then trained a classifier to annotate simplified behaviors including rest, turn, and cruise. For simplicity, we defined a cruise as a forward swim with at least five measured extrema, or approximately three body oscillations.

**Figure 2.**
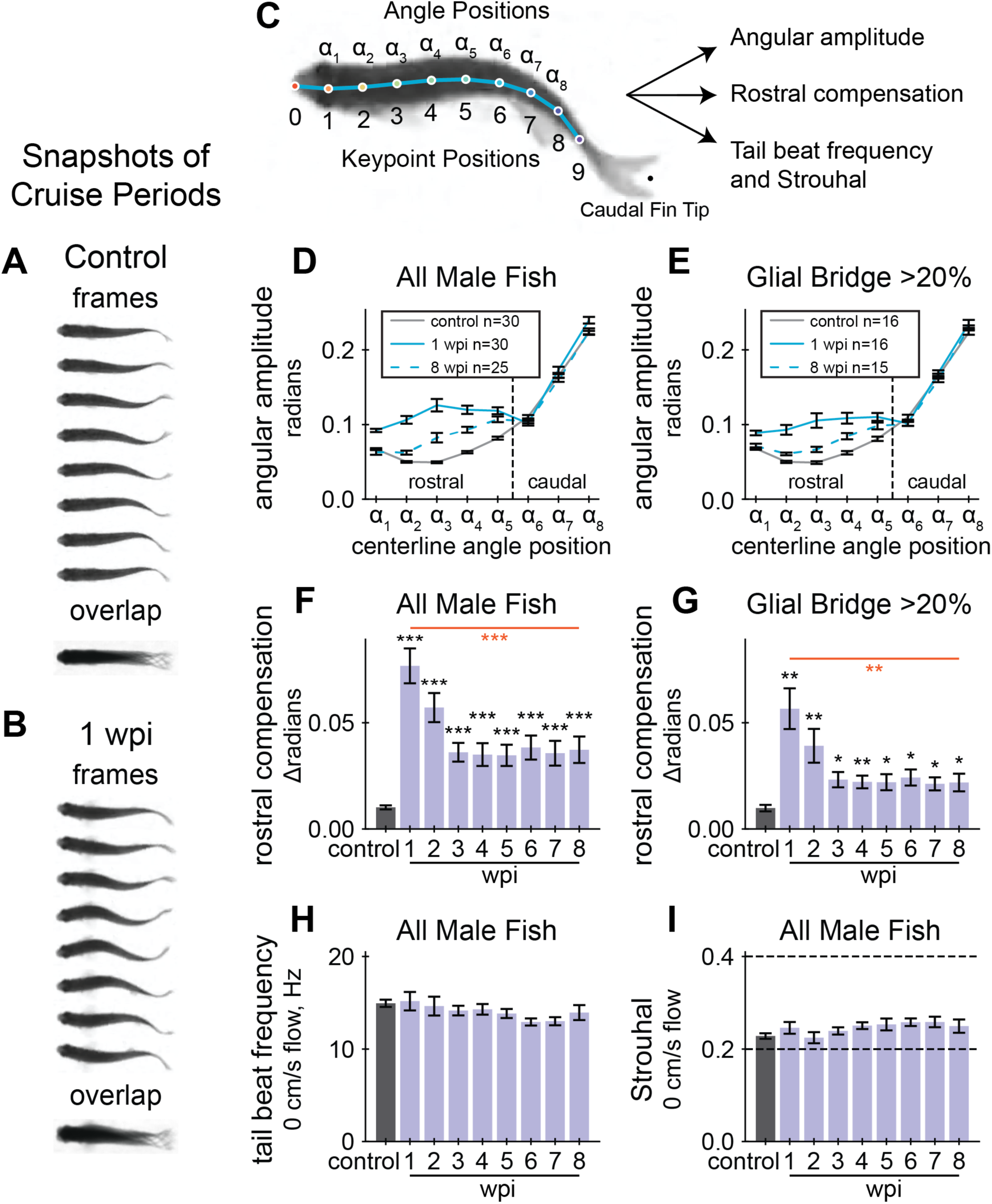
Cruise quality quantifications during SC regeneration. **(A, B)** Cruise periods at a water current velocity of 10 cm/s. Snapshots of swim behavior prior to injury are shown in A and at 1 wpi in B. Overlayed swim forms shown below the separate frames highlight defective caudal movement after SC transection **(C)** Experimental schematic to develop quantifiable gait quality measurements. A diagram showing keypoint and angle positions labeled on a fish’s centerline. A DeepLabCut model was trained to identify the keypoints, then the curvature between three adjacent keypoints were calculated at each angle position. The rostral region of the fish corresponds to angle positions ⍺_1_ to ⍺_5_. **(D)** Cruise curvature profiles for male fish before injury, acutely injured fish at 1 wpi, and recovered fish at 8 wpi. Each cruise curvature profile represents the mean lateral angular amplitude along the dorsal centerline while cruising. The vertical dotted line separates rostral and caudal positions, demonstrating that acutely injured fish swim with markedly elevated curvature in the rostral portion of their body. **(E)** Cruise curvature profiles (control, 1wpi, and 8wpi) for male fish that regenerated more than 20% of the glial tissue at the lesion site. **(F)** Quantification of rostral compensation in male control fish prior to SC injury and at 1 to 8 wpi. Rostral compensation represents the displacement between cruise profiles. This score is the maximum distance, on the vertical axis, between rostral positions of a cruise profile from the control profile. Angle positions ⍺_1_ to ⍺_5_ from Panel D were used to define the rostral region. **(G)** Rostral compensation scores for male fish that regenerated more than 20% of the glial tissue at the lesion site. **(H)** Tail beat frequency for male fish in still water (0 cm/s), measured at each assayed week. ANOVA was not significant for tail beat frequencies. **(I)** Strouhal numbers for male fish in still water (0 cm/s), measured at each assayed week. Strouhal number is a unitless value related to vortex shedding mechanics and is defined as tail beat frequency times peak-to-peak amplitude of the tail tip divided by speed of forward motion. ANOVA was not significant for Strouhal numbers. Error bars depict SEM and statistical significance was determined by one-way repeated mixed measures ANOVA followed by post-hoc Student’s T-tests. p-value markers in black represent comparisons between each time point post-injury relative to control measurements prior to injury. Red horizontal bars and p-values marked in red show significance between 1 wpi and 8 wpi. ***P<0.001; **P<0.01; *P<0.05; ns, P>0.05.

Our first approach to probe swim gait was to examine posture variation within healthy and injured animals without regard to behavior. To this end, we used principal components analysis (PCA) to decompose all poses where the fish was not near a wall. PCA was performed for all fish and all assays (Figure S2A), separately for males (Figure S2B) and females (Figure S2C), and at control and recovery endpoints 1 and 8 wpi per sex group (Figure S2D-E). Exploring the principal components of control poses, merely three components were required to explain as much as 98% of the variance in both groups. Using poses from 1 wpi, four components were required to capture 98% of the variance. At the end of 8 weeks in recovery, as many as five components were required to capture 98% of the variance. These results indicated that centerline posture increased in complexity after injury and through recovery.

Next, using cruise episodes annotated by the machine learning classifier, we measured body curvature amplitudes and frequencies at each measured angle position. Curvature at each angle position was calculated using the corresponding keypoint position and two adjacent keypoint positions (Figure 2C). We measured center-to-peak (half of peak-to-peak) curvature amplitude at all angle positions during cruise oscillations. Cruise amplitudes along the centerline were plotted for control, 1 and 8 wpi (Figure 2D). At 1 wpi, we observed a significant change in angular amplitude at angle positions 1 to 5 compared to uninjured controls. Cruise amplitudes appeared closer to control values at 8 wpi. These findings indicate that acutely injured fish compensate for paralysis using the rostral portion of their body and that compensation is less pronounced after cellular regeneration. Hypothesizing that fish swim more naturally if they regenerated well, we separately plotted cruise amplitudes for fish that regenerated at least 20% of their glial tissue at the lesion core (Figure 2E). Well-regenerated fish appeared to swim using amplitudes closer to control values. These results were also observed for female fish, although fewer female fish, 5 females versus 16 males, recovered more than 20% of their glial tissue at the lesion (Figure S3B). We quantified maximal rostral compensation and noted that this feature of swim gait does recover toward control levels (Figure 2F). We also measured summed rostral amplitude (Figure S4D). Fish with more regenerated SC tissue recovered healthier gait on average, indicated by lower rostral compensation at 8 wpi (Figure 2G). Individual recoveries were more pronounced: 17 fish at 8 wpi exhibited rostral compensation lower than 3 standard deviations above the control mean, and these fish had high average cellular regeneration: 59% glial bridging, 57% proximal axon regrowth, and 34% distal axon regrowth. All fish with compensation score lower than 0.025 (2.49 standard deviations above the control mean) had at least 20% glial bridge. These results indicate that rostral compensation is most pronounced during earlier stages of SC regeneration and that stronger cellular regeneration is associated with healthier swim quality.

In addition to angular amplitude, we also measured cruise frequency, defined as the number of complete oscillations at each angle position per second. Regardless of injury, the rate of movement during cruises was nearly uniform along the fish’s body (Figure S4A). Like amplitude, average body cruise frequency was also disrupted by injury and improved over time (Figure S4B-C). Although functional abilities appeared different between the sexes according to established functional metrics, there was no evidence that rostral compensation differs between males and females at 1 wpi (Student’s t-test p-value = 0.6), suggesting that SC transection affects the gait of both sexes equally and independently of swim capacity differences.

Cruising zebrafish produce thrust by ejecting vortices of water behind them (28). Strouhal number (St) is a unitless value related to vortex shedding mechanics and is defined as tail beat frequency times peak-to-peak amplitude of the tail tip divided by speed of forward motion. Cross species research has concluded that St between 0.2 and 0.4 are most physiologically efficient (38). We calculated tail beat frequency for fish swimming in static water (0 cm/s flow velocity) and did not observe a significant change after injury (Figure 2H). Using displacement amplitude of the caudal fin, fin flick frequency, and swim speed during cruises, we calculated St for all cruises during the first five minutes (0 cm/s flow velocity) of the swim assay. Average St was within or near the efficiency limits throughout the experiment (Figure 2I, Figure S3F). These results suggest SCI does not preclude efficient swim thrust, likely due to rostral compensation during periods of active cruising.

### Injury-induced lateral scoliosis is irreversible by 2 weeks post-injury

Before exploring centerline-derived gait more comprehensively, we had to address injury-induced scoliosis. Although all fish in the experiment appeared healthy before injury, some fish developed extreme body curvature after injury. We quantified scoliosis as the summed curvature at eight angle positions along the fish’s dorsal centerline (Figure 2C) averaged over the entire swim. Representative fish with their scoliosis scores are presented in Figure 3A. Our analysis indicated that of the 30 male and 30 female fish, 4, 2, and 1 male and 4, 3, and 2 female fish presented scoliosis scores >1.34, >2.77, and >3.28, respectively. Pairwise comparisons of scoliosis scores between time points, using Spearman correlation, showed that relative scoliosis rankings stabilized after 2 wpi (Figure 3B). Fish did not exhibit noticeable scoliosis at 1 wpi, nor did scoliosis scores at 1 wpi correlate with the following weeks. Scores at pre-injury and 1 wpi weakly correlated with subsequent time points since the fish were all straight bodied at the beginning of the experiment and had not developed scoliosis by 1 wpi. The randomness of scoliosis rankings at 1 wpi and stability of rankings after 2 wpi are presented in Figure 3C using trendlines for each fish colored by the fish’s rank at a specific point in recovery. As an alternative, unsupervised posture-driven recovery measurement, we used a nearest-neighbors approach to quantify postural abnormality for each assay. Taken as a ratio compared to representative posture observed in preinjury assays, we called this measurement “posture novelty”. Posture novelty was closely associated with scoliosis scores. Pairwise rank correlations of scoliosis scores suggest that injury-induced scoliosis is irreversible by 2 wpi, and these results altogether revealed that scoliosis is an important factor in the variation of poses observed in the experiment at 8 wpi (Figure S2D-E).

**Figure 3.**
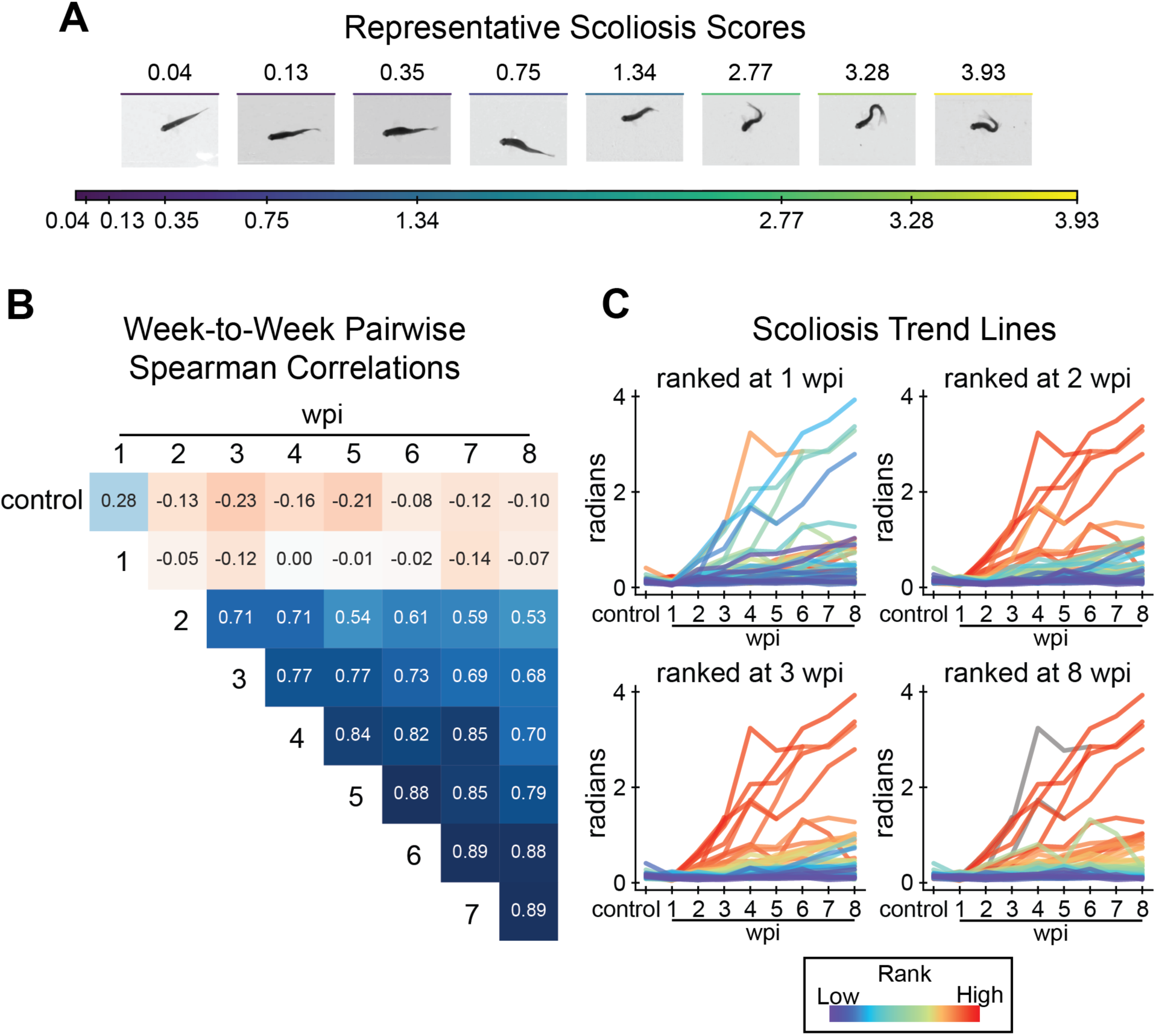
Assessment of lateral scoliosis among all fish, male and female, in the tracking experiment. **(A)** Visualization of the scoliosis score. The scoliosis score is the sum of curvature measured at angle positions on the centerline at rest (see methods). To demonstrate how this score relates to body curvature, frames were taken from representative assays identified using k-medoids clustering of scoliosis scores of all assays. **(B)** Pairwise comparison of scoliosis scores between weeks by Spearman’s rank correlation. Scoliosis scores correlate weakly between control and 1 wpi to other assays. Scores are most strongly correlated from 2 wpi onward. **(C)** A demonstration that scoliosis ranks stabilize after 2 wpi. Scoliosis trend lines for individual fish are plotted over all assays (x-axis), colored by rank at 1, 2, 3, and 8 wpi (top-left to bottom-right). Fish scoliosis trend lines are colored on a rainbow scale from red (most severe) to violet (least), and colored gray if the fish did not survive to the ranked week.

### Gait recovery plateaus around 3 weeks post-injury

Because our rostral compensation score is just one feature of complex locomotion, we next set out to comprehensively analyze cruise gate recovery by embedding cruise episodes into a latent space. To reduce the effects of body shape on gait, we omitted fish with a scoliosis score greater than 0.35 at 8 wpi. For computational efficiency, we performed embedding on representative cruise episodes, exemplars, sampled from each assay. To explore the data in posture space, we decomposed poses using principal components analysis. Cruise traces in posture space manifested as ellipses, similar to previously described traces for larval zebrafish (20, 21). We noted that 1 wpi cruise traces are both distinct from preinjury cruises and visually similar for different animals within the same time point (Figure 4A). Then, we embedded cruise traces using UMAP for three-dimensional visualization. We plotted cruises from highly regenerative fish, those that recovered at least 20% glial tissue at the lesion (Figure 4B), separately from cruises of fish with relatively poor cellular regeneration (Figure 4C). To interpret this embedding, average UMAP values at each time point for decomposed cruise episode exemplars were plotted for highly (Figure 4D) and poorly (Figure 4F) regenerating fish. For both fish groups, average UMAP values exhibited the most change by 3 wpi. Similarly, for both highly and poorly regenerating fish, rostral compensation recovery slowed around 3 wpi (Figure 2F,G; Figure S4B,D). Kernel density estimates were calculated for each time point in each UMAP dimension, separated by cellular regeneration outcome (Figure 4E,G). According to the UMAP kernel density estimates, 8 wpi cruises did not inhabit the same embedding space as preinjury cruises. Cruises for female fish were embedded in the same space and the results were visually similar (Figure S5A-G). This temporal cruise embedding suggested that gait recovers to the greatest extent by 3 wpi and that injury-induced gait changes are incompletely resolved at 8 wpi.

**Figure 4.**
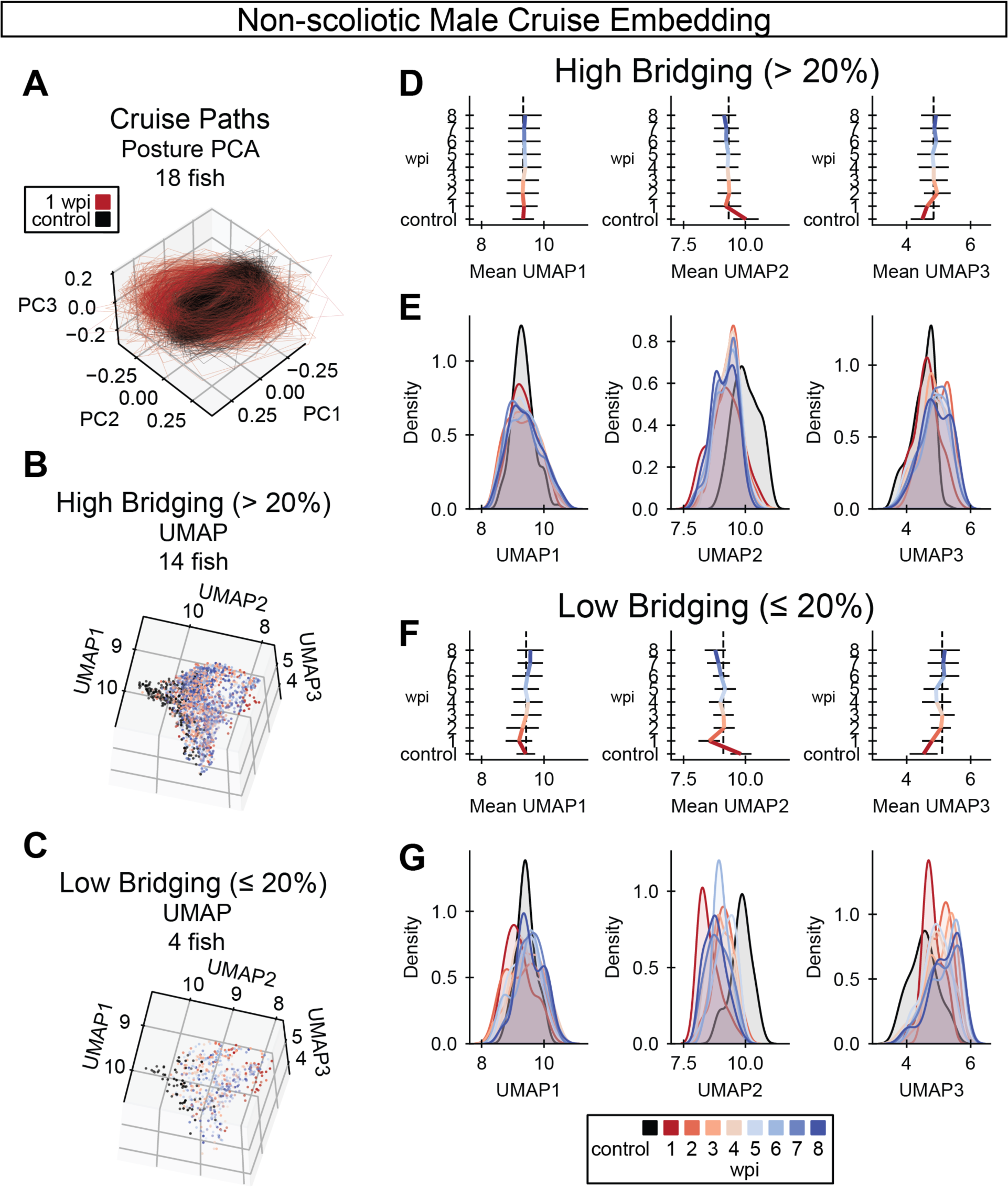
UMAP embedding of cruise behaviors during SC regeneration. **(A)** Cruise poses from control (black) and 1 wpi (red) male zebrafish were decomposed using principal component analysis (PCA). Male fish with scoliosis score < 0.35 at 8 wpi are shown. In this posture space, cruise trajectories from acutely injured fish at 1 wpi are markedly distinct from control fish trajectories. **(B-C)** Cruise poses were embedded using UMAP with precomputed dynamic time warping (DTW) distances. Cruise poses were analyzed for fish that regenerated well (glial bridging > 20%) in B and for fish with compromised cellular regeneration (glial bridging < 20%) in C. **(D)** Gait recovery trendlines in UMAP 1, 2, and 3 (left to right) for fish that regenerated well. Colored bars indicate the mean trend of each UMAP component. Error bars are standard deviations. The vertical dotted black lines align with the average UMAP value at 3 wpi to indicate that most change in each UMAP occurs before 3 wpi. **(E)** Gaussian kernel density estimates for the regions of UMAP space occupied by cruises at each week post-injury for fish that regenerated well. The x-axes are matched between Panels D and E for comparison. **(F)** Gait recovery trendlines in UMAP 1, 2, and 3 (left to right) for fish with compromised cellular regeneration. Colored bars indicate the mean trend of each UMAP component. Error bars are standard deviations. The vertical dotted black lines align with the average UMAP value at 3 wpi to indicate that most change in each UMAP occurs before 3 wpi. **(G)** Gaussian kernel density estimates for the regions of UMAP space occupied by cruises at each week post-injury for fish with compromised cellular regeneration. The x-axes are matched between Panels F and G for comparison.

### Selected functional measurements correlate with cellular repair at 8 weeks post-injury

We next explored relationships between the full array of functional and cellular regeneration metrics at 8 wpi. We measured Spearman correlations between all 8 wpi measurements, pairwise, then performed hierarchical clustering on the correlation matrix (Figure 5A). While not all measurements correlated well with cellular regeneration metrics, some did, including quantifications of capacity, behavior, and gait. Selected measurements of these classes were plotted against glial bridging: swim distance, a swim capacity measurement (Figure 5B); burst frequency quantified using active posture, a behavior measurement (Figure 5C); and rostral compensation, a gait quality measurement (Figure 5D). Percent glial bridging correlated strongly (r_s_ > 0.6) with both proximal and distal axon regrowth (Figure 5E). Assays could only be included in the heatmap if the fish exhibited every measurable attribute, and because two fish did not cruise in the 8 wpi swim assay, their data was omitted from the heatmap in Figure 5A. There were also three male and three female fish that did not survive to the end of the experiment. Hence correlations differ slightly between Figure 5A and Figure 5B-E.

**Figure 5.**
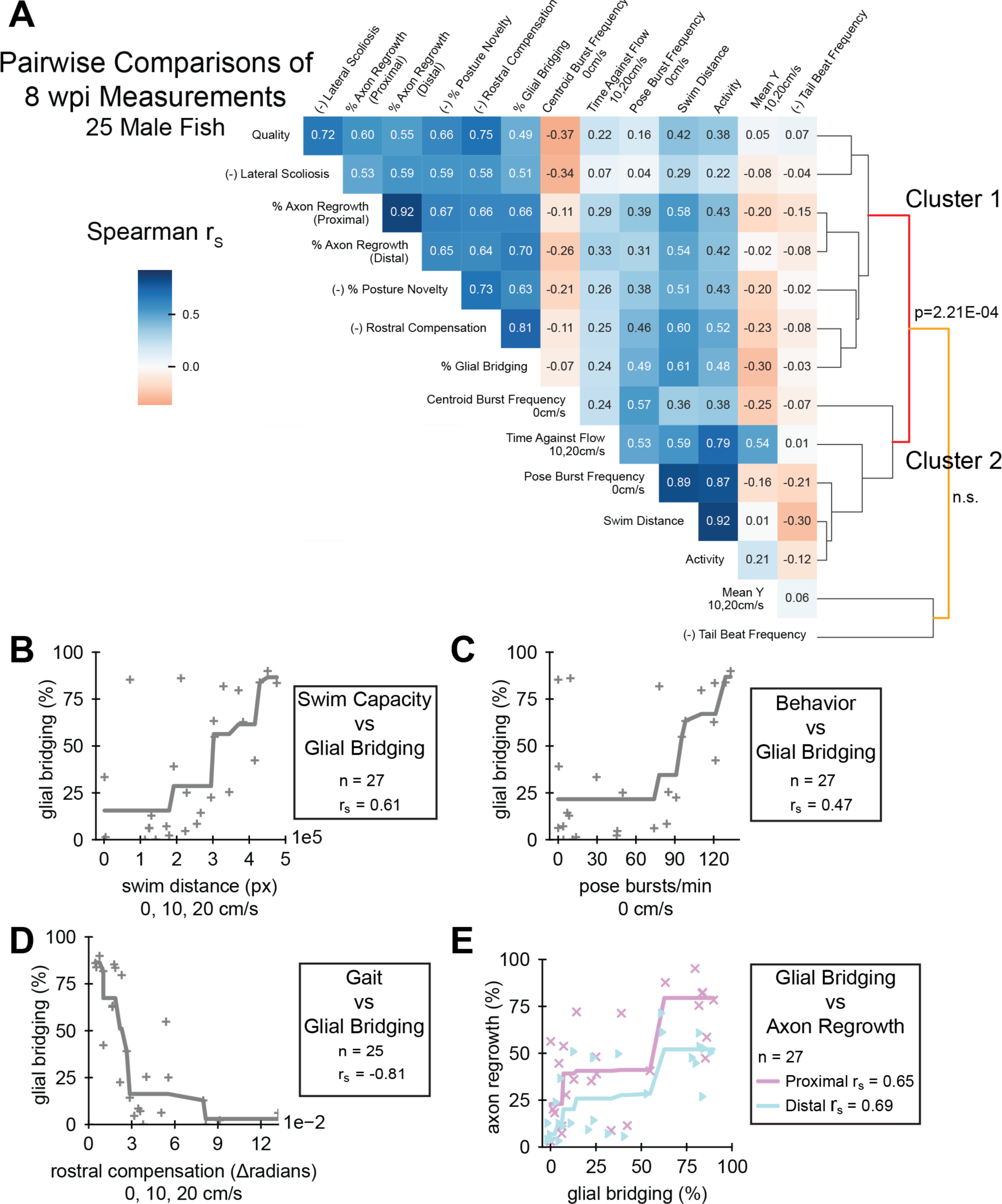
Pairwise comparisons of spinal cord regeneration metrics at 8 wpi. **(A)** Heatmap of Spearman’s rank correlations (r_s_) between the cellular and functional measurements taken. Regeneration metrics include swim capacity, swim quality, structural, and neurological measurements. A dendrogram representing similarities between correlation patterns of measurements is shown. Male fish at 8 wpi were analyzed. Fish that could not be measured in all attributes were omitted. Correlations are reported regardless of p-value. For the sake of meaningful clustering, a measured attribute was multiplied by -1 if its average was increased at 1 wpi compared to controls. These attributes are marked with the prefix “(-)” on the label. **(B-D)** One metric each of swim capacity (swim distance), behavior (burst frequency), and gait (rostral compensation) were plotted against glial bridging to demonstrate their correlation. Because the Spearman correlation operates on ranked values, we fit monotonic splines to plots where r_s_ were significant (p < 0.05) to visualize possible associations. **(E)** Scatter plot showing a strong association between the size of regenerated glial tissue and axon regrowth, proximal and distal.

Comparing pairwise correlations at 8 wpi, swim capacity and behavior measurements generally correlated strongly with one another and resided in a swim capacity associated cluster that we labeled Cluster 2 (Figure 5A). Such measurements include distance, activity, pose burst frequency, and time swimming against the flow. Mean y-position correlated moderately with time swimming against the flow, but weakly with the other swim capacity measurements. Most, but not all, capacity measurements were at least moderately (r_s_ > 0.3) correlated with glial bridging and axon regrowth. At 8 wpi, tail beat frequency, which did not change significantly after injury or through recovery (Figure 2H, Figure S3E), did not correlate well with any other functional, structural, or cellular measurement. A separate cellular regeneration associated cluster, labeled Cluster 1, contained all cellular regeneration measurements, rostral compensation, perceived swim quality, scoliosis, and posture novelty. Unlike the swim capacity associated cluster, all metrics in Cluster 1, were at least moderately, positively correlated with each other. Of all functional metrics, rostral compensation correlated most strongly with perceived quality and percent glial bridging, and with axon regrowth to the same extent as posture novelty. Together, this analysis showed that not all the swim capacity metrics we measured were representative of the SC’s condition at 8 wpi, and that rostral compensation is a neurologically associated measurement of gait quality.

As previously described, female fish in this experiment regenerated more poorly than male fish. Nonetheless, a similar clustering of 8 wpi measurements from female fish yielded the same clusters (Figure S6A). Spearman rank correlation of swim distance, pose burst frequency, rostral compensation, and axon regrowth with glial bridging did not reach statistical significance for female fish (Figure S6B-E). Despite their non-significant p-values, rostral compensation and proximal axon regrowth showed the highest correlation values with glial bridging (Figure S6A). These results support our model that rostral compensation is the most neurologically relevant of all metrics in this tracking study.

### Tensor decomposition of all assay measurements corresponds with outcomes at 8 wpi

We applied tensor component analysis (TCA) to explore recovery trajectories. To prepare the data, we concatenated measurements from all 9 assays of all 60 fish into a rank-3 tensor. The software that we used for TCA did not allow missing values, and we decided not to use interpolation, so 16 fish (6 male, 10 female) had to be omitted. We selected parameters for the model by optimizing for low model error and high (>0.8) model similarity (Figure 6A). After selecting parameters for the model, we obtained 7 tensor components including factors for each axis: fish, assay, and functional measurement (Figure 6D). To visualize factors, we sorted fish by the size of their glial bridge at 8 wpi and sorted assays temporally. By manual inspection, values for many fish factors showed a directional trend, implying that the trajectories captured in the factor values represented a functional output relevant to neurological recovery. We explored this relationship further by clustering fish according to their factor values. We selected a distance threshold parameter that yielded the most clusters given a high minimum allowed cluster size (Figure 6B). The five resulting clusters of fish were roughly the same size, including between 7 and 11 fish in each cluster. Cluster averages for measurements taken at 8 wpi were plotted in a heatmap, with clusters sorted by average glial bridging (Figure 6C). When ordered by average glial bridging in the cluster, clusters also sorted according to average scoliosis, posture novelty, perceived quality, and rostral compensation. In the same ordering but omitting cluster 1, clusters 2 to 5 also sorted according to average measurements of axon regrowth (proximal and distal), swim distance, activity, time against flow, posture burst frequency, and mean y (but in the opposite direction than expected). The only measurement averages that were not sorted in this ordering were centroid burst frequency and tail beat frequency. Fish factor values in Figure 6D were colored by the fish’s cluster according to the cluster’s average glial bridging. We expected that assay factors would exhibit a displacement-to-recovery trend from 1 to 8 wpi, mimicking the recovery trend of most functional metrics (Figure 1B-G, Figure 2F). Although some assay factors did capture this trend, many diverged away from control through recovery, and the values of TC1 and TC2 did not change dramatically from control until 2 wpi, capturing a trend more similar to the scoliosis score (Figure 3C). We interpret the diversity of assay factor trendlines to represent complexities in the differing biological signals of the functional measurements and longitudinal support for non-uniformity of 8 wpi metric correlation signals shown in Figure 5A. Overall, tensor component analysis and fish factor clustering indicate that recovery trajectories correspond strongly with 8 wpi cellular regeneration measurements.

**Figure 6.**
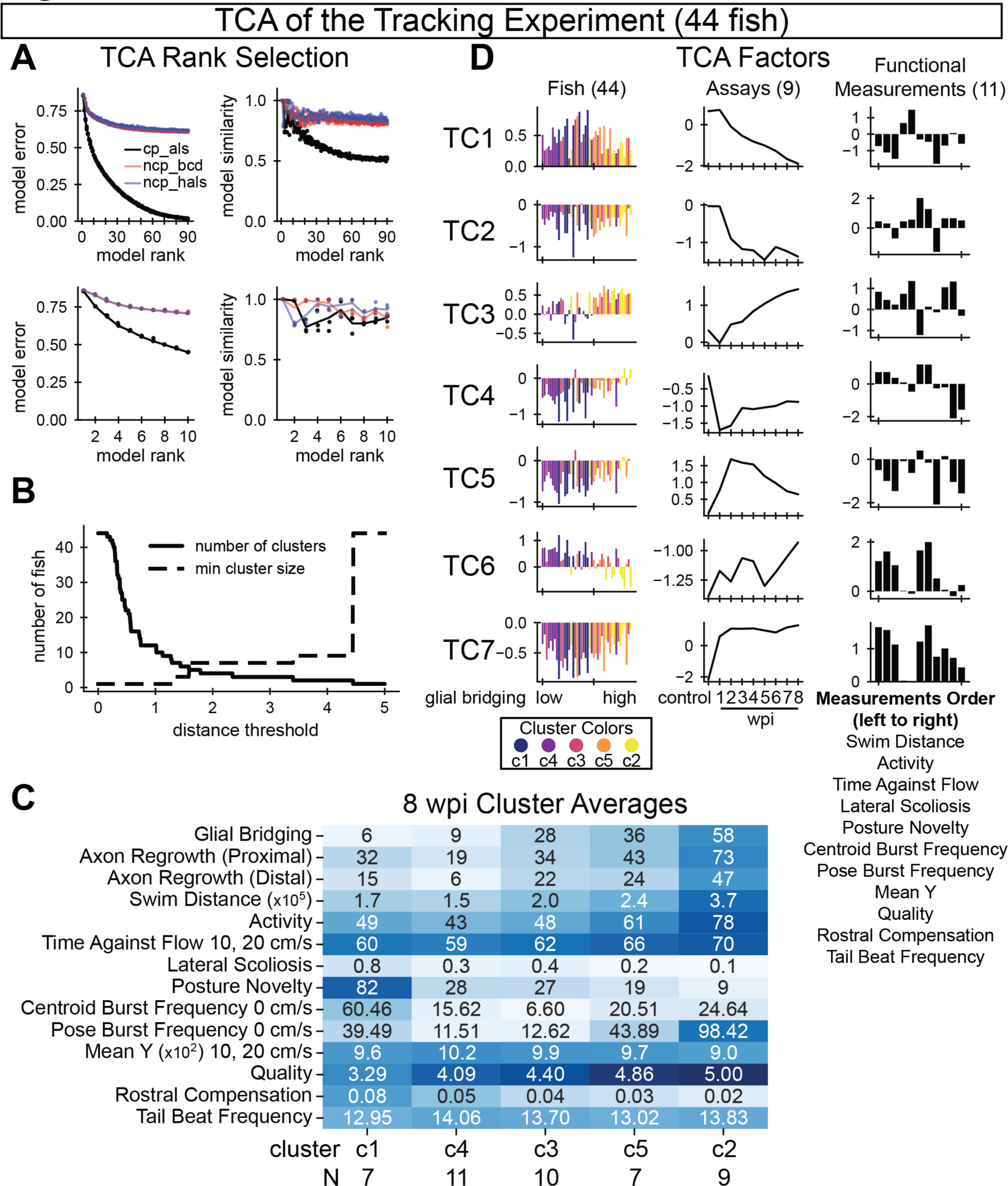
Tensor component analysis of swim behavior during SC regeneration. **(A)** Model selection parameters. Plots on the left show model error various ranks (x-axis). Plots on the right show model similarity for various ranks (x-axis). Unlike PCA, TCA is not deterministic. Thus, multiple models were trained and compared. Three model types are shown. To achieve an optimal balance between low model error and high model similarity, we chose a rank-7 canonical polyadic decomposition by alternating least squares (cp_als). **(B)** Selecting an appropriate distance threshold for clustering. We clustered fish according to their TCA factors using agglomerative clustering. To maximize the number of clusters while keeping a high minimum cluster size, we chose 1.7 as our distance threshold with five clusters identified. **(C)** Recovery outcomes for the five identified fish clusters, sorted by average glial bridging, low to high, from left to right. Each row’s color is normalized such that the darkest blue corresponds to the maximum observation among all fish for that measurement. **(D)** Factors from a rank 7 tensor decomposition corresponding to each axis of the full data tensor: fish, assays (wpi), and functional measurements. Fish are sorted on the x-axis by their 8 wpi measured percent glial bridging and colored according to the rank of the cluster with respect to average glial bridging. Assays are sorted temporally. Measurements are listed below the plots in descending order. For panels A-D, 44 out of 60 total fish were analyzed. 16 fish were omitted either due to early death (6 fish) or because cruise waveform statistics were insufficient for at least one assay (10 fish).

### Swim function at 2 weeks post-injury is predictive of cellular regeneration at 8 weeks

Swim distance and rostral compensation were two functional measurements that most correlated with SC regeneration at 8 wpi. Swim distance and rostral compensation measure different functional classes: swim capacity and gait quality, respectively. They also clustered separately in the pairwise metric analysis and recovered at different rates. Thus, as distinct functional outputs that correlate well with cellular regeneration, we hypothesized that a combination of swim distance and rostral compensation assessed during early stages of SC regeneration could predict injury outcomes at 8 wpi. To test this hypothesis, we averaged 1 and 2 wpi distance and rostral compensation measurements and plotted them against one another colored by 8 wpi glial bridging (Figure 7A). As expected from the correlation analysis, fish with high glial bridging generally swam longer distances and with lower rostral compensation. We ranked fish according to swim distance and negative rostral compensation, then added the two ranks to create a wellness score for early recovery. We divided male and female data separately into two equally sized groups across each sex’s wellness score median: a highly regenerative prediction group and a poorly regenerative one. Both percent glial bridging and proximal axon regrowth were significantly higher in the highly regenerative group compared to the poorly regenerative group (Figure 7B). Since this analysis was performed after our tracking experiment was completed and final outcomes were measured, a new experiment was performed to validate the prediction method.

**Figure 7.**
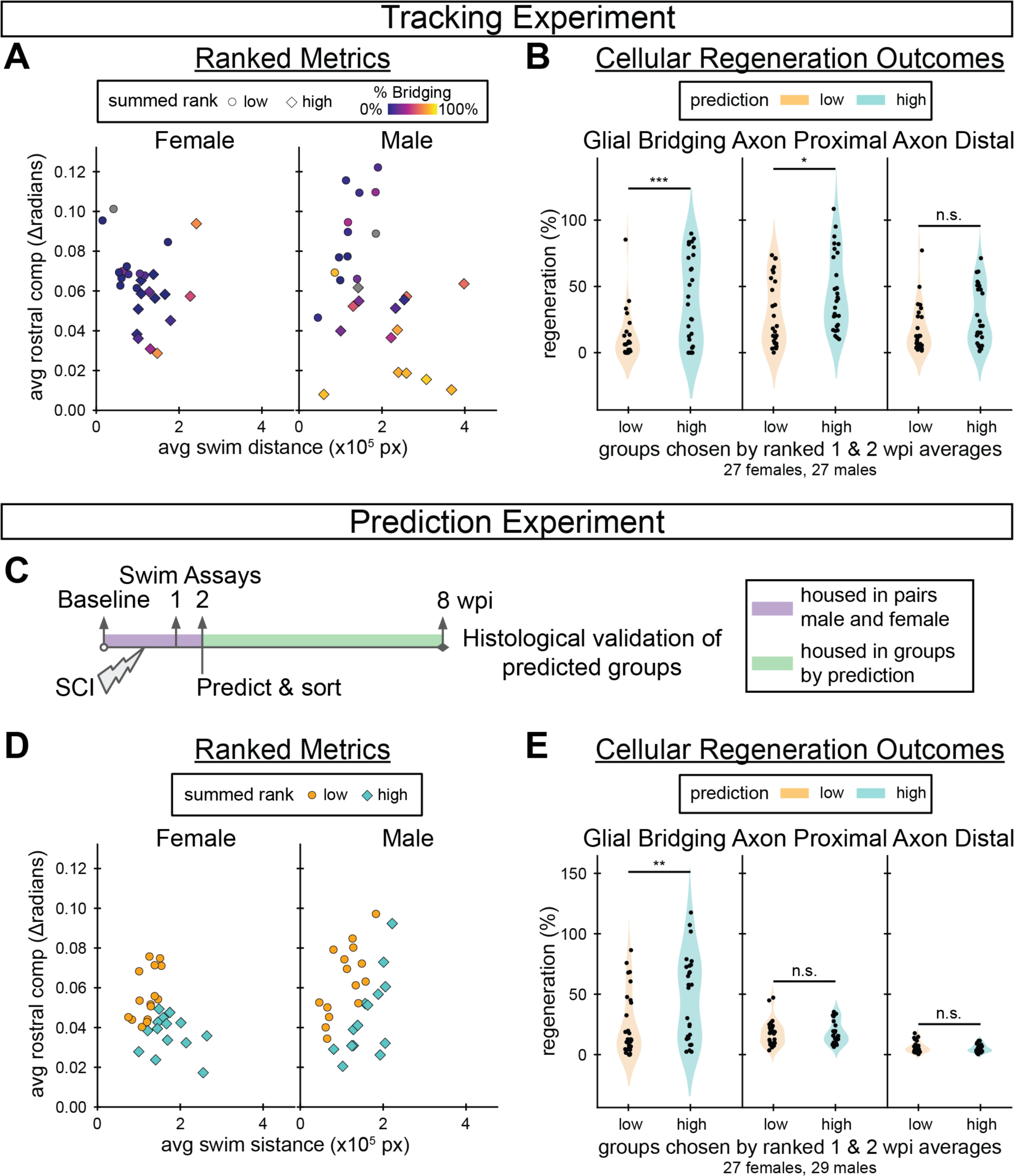
Predicting 8 wpi regeneration outcomes using measurements taken by 2 wpi. **(A)** Tracking experiment: swim distance versus rostral compensation was averaged between 1 and 2 wpi. Dots represent individual fish and were colored based on the extent of glial bridging measured at 8 wpi for each fish. Fish that would develop larger glial bridges tended to swim further and exhibit less rostral compensation, in lower right side of each graph. Each fish was assigned a prediction score by adding its rank according to average swim distance to its rank according to average negative rostral compensation, relative to its sex group. **(B)** Tracking experiment: 8 wpi glial bridging and axon regrowth outcomes for groups ranked high in swim distance and low in rostral compensation between 1 and 2 wpi (“high”) compared to those ranked low in swim distance and high in rostral compensation between 1 and 2 wpi (“low”). **(C)** Schematic overview of the prediction experiment. A total of 60 adult fish (30 males and 30 females) were housed in male-female pairs to track their identities until low/high regeneration predictions were assigned, then housed in prediction groups. At 8 wpi individuals were separated again to correlate final measurements from the endurance and behavior assays with cellular regeneration. Behavior was assayed before injury, 1 wpi, 2 wpi, and 8 wpi. Predictions were made, as described, after the 2 wpi behavior assay. **(D)** Prediction experiment: swim distance versus rostral compensation was averaged between 1 and 2 wpi. Dots represent individual fish and were colored based on the predicted outcome. **(E)** Prediction experiment: 8 wpi glial bridging and axon regrowth outcomes for each prediction group. Statistical significance in panels B and E was determined by Student’s t-test. p-values represent comparisons between prediction groups. ***P<0.001; **P<0.01; *P<0.05; ns, P>0.05.

In the prediction experiment, we predicted regeneration outcomes for sixty (30 male, 30 female) fish using the same wellness score described above (Figure 7C). Swim behavior was assayed before injury in addition to 1, 2, and 8 wpi. Fish were housed in male-female pairs to track their identities until swim assays were performed at 2 wpi. Predictions were made separately for males and females, then fish of both sexes were mixed randomly between tanks within the prediction groups, 5–7 fish per tank. Predictions were made using the median wellness scores in the prediction experiment to divide the groups (Figure 7D). At 8 wpi, glial bridging was significantly elevated in the highly regenerative prediction group compared to the poorly regenerative prediction group (Figure 7E). The results of this prediction experiment indicated that it is possible to predict 8 wpi cellular regeneration outcomes from a subset of 2 wpi measurements.

We clustered the 8 wpi functional and regeneration measurements in the same way as we did for the tracking experiment. In doing so, we discovered that endurance, a measure of swim capacity that was not evaluated at any other timepoint or in the tracking experiment, was a member of the neural health-associated cluster, Cluster 1, not the swim capacity cluster (Figure S7A). Unlike the tracking experiment, swim distance and pose burst frequency were not associated with bridging to the same extent (Figure S7B-C), and axon regeneration was measured lower than expected for all fish (Figure S7E). Rostral compensation, however, still correlated more strongly with glial bridging than any other measurement (Figure S7D). These findings provided additional evidence that rostral compensation is a neurologically associated gait quality measurement, and suggested that results derived from gait quality replicate more reliably than those derived from swim capacity.

## Discussion

This study explores connections between functional and cellular regeneration measurements during SC regeneration in adult zebrafish. Fish were individually housed to enable their longitudinal tracking for 9 weeks. In comparison with alternative methods of tracking fish identities such as fin clipping, dye injection, passive integrated transponder tagging or skin pattern recognition, physical separation was the method of choice to avoid introducing additional injuries or imaging steps to our experimental pipeline (39). It is important to note that, compared to the well-known elevated regenerative capacity of zebrafish SC tissues, regeneration efficiency was reduced in this cohort of 60 fish. Although it remains to be determined whether physical separation negatively impacts SC regeneration, we propose that our previously established injury and post-operative standards, including group housing, support optimal regeneration conditions (10, 40). Even in such conditions where functional recovery is expected to continue to 8 wpi, some fish experience negligible functional improvement after 6 wpi (3). However, our purpose was not to demonstrate SC regenerative capacity in zebrafish, but to compare traditional and new functional measurements and their associations with structural recovery. Our fish exhibited significant recovery up to 6 wpi, glial and axon bridging were highly correlated, and all three measurements of cellular regeneration showed similar correlation patterns with examined functional and structural metrics. Although functional recovery appeared different between the sexes, we saw no evidence that the cruise gait of males and females are affected differently by SC transection, comparing male versus female rostral compensation scores at 1 wpi using Student’s t-test, (p-value = 0.6). Thus, irrespective of whether housing conditions for this study were detrimental to recovery, the broad range of regeneration observed in this study was not only well suited, but also retrospectively ideally tailored, for our purposes. Our tracking experiment allowed us to perform comprehensive correlative analyses of recovery metrics over a wide range of regeneration parameters.

Our study introduces rostral compensation score as a scalable quantification for gait quality. In both the tracking and prediction experiments, rostral compensation correlated more highly with glial bridging than any other measurement at 8 wpi. The strong correlations we observed between rostral compensation and cellular regeneration measurements support a model in which rostral compensation score reflects the compensatory effort paralyzed fish make to produce sufficient thrust to move. Whether fish optimize their Strouhal number by learning to adjust the appropriate amount of rostral compensation or whether the optimal Strouhal number is physiologically enforced by the natural elasticity of a fish’s body remains to be explored (41). The calculation of rostral compensation was entirely automated after training a classifier to annotate cruise behaviors from centerline posture. As a recovery metric in the tracking experiment, average rostral compensation scores showed 50% recovery at 8 wpi. Notable and significant changes in rostral compensation occurred by 3 wpi, suggesting a threshold of cellular regeneration is required for healthy functional outcomes and that cellular regeneration beyond 3 wpi has a diminished effect on functional recovery. Overall, our findings show that rostral compensation is a neural health-associated measurement that recovers earlier than swim capacity.

To achieve a more comprehensive analysis of cruise gait recovery beyond single gait features such as rostral compensation, cruise episodes were embedded with UMAP using dynamic time warping distances. Surprisingly, even after selecting for fish that showed elevated regeneration metrics, 8 wpi cruises showed a clearly distinct distribution compared to control cruises prior to injury. These findings suggest that some injury-induced gait feature persists for many fish after apparently successful recovery. When an abnormal swim gait develops during recovery, it could be due to mistaken axon connections or suboptimal tissue remodeling. Specific SC neurons and regions are known to be associated with aspects of zebrafish locomotion (42, 43). After SC transection, as axons reconnect, fast and slow V2a interneurons regrow axons along the rostro-caudal axis, connecting to neurons of the same type and enabling specific features of functional recovery (12). In addition to axon regrowth, it is estimated that as much as a third of SC tissue shows evidence of cell division during SC regeneration (44). Moreover, recent mammalian studies indicate that certain neurons required to restore walking after SC injury are not the same as the neurons used for the same behavior before injury (45). Together, these various modes of cellular regeneration may explain why, given the natural variation in recovery outcomes, the kernel density estimates of UMAP-embedded cruises did not completely overlap with those of uninjured control cruises. We propose that differential swim gait development during the recovery process could be due to misguided axon regrowth or suboptimal tissue remodeling, and that further cellular and behavioral regeneration studies are needed to better understand the cellular basis of swim recovery in zebrafish.

We performed a separate experiment to explore the hypothesis that rostral compensation and swim distance measurements at 1 and 2 wpi can be used to predict 8 wpi cellular regeneration outcomes. The prediction experiment, while not completely successful at predicting significantly different outcomes for both glial bridging and axon regrowth, nonetheless provides a proof of concept for future work. In both the tracking and prediction experiments, glial regeneration outcomes were significantly different, which indicates that early functional measurements can be predictive of cellular regeneration measured six weeks later. Moreover, our prediction experiment provides validation for rostral compensation as a neurologically meaningful functional metric, and evidence that swim endurance, which was not evaluated in our tracking experiment, is also associated with neurological health. Importantly, both the tracking and prediction experiments support our conclusion that rostral compensation provides the strongest correlation with cellular regeneration than any other functional measurement that we examined. Comparing the two experiments, we likely overestimated swim distance’s predictive power, and knowing that gait recovers relatively early, rostral compensation alone may be the best metric for outcome prediction, which remains to be tested.

## Conclusion

Our study supports standard functional analysis, demonstrating that recovered swim capacity measurements correlate with spinal cord regeneration. We introduce rostral compensation as a gait-driven measurement that correlates more strongly with and likely provides a more accurate functional proxy for spinal cord regeneration than existing functional regeneration metrics. Through posture analysis, we discovered that the fish that would develop injury-induced lateral scoliosis had a set trajectory by two weeks following SC transection. We found that, in still water, fish swam in a mechanically efficient manner both before and after injury. Finally, we discovered that grouping fish according to their rank in functional attributes two weeks into recovery can divide the fish into groups that have statistically divergent cellular regeneration outcomes at eight weeks post-injury. We propose that developing a gait-driven prediction algorithm could offer powerful benefits to the zebrafish research community such as non-invasively tracking musculoskeletal wellness or reducing the time required to complete SC regeneration experiments.

## Acknowledgments

We thank T. Darveniza, J. Jensen and Z. Pincus for discussion and the Washington University Zebrafish Shared Resource for animal care. This research was supported by grants from the NIH (R01 NS113915 and R01 NS123708 to MHM).

## Supplementary Figures

**Figure S1.**
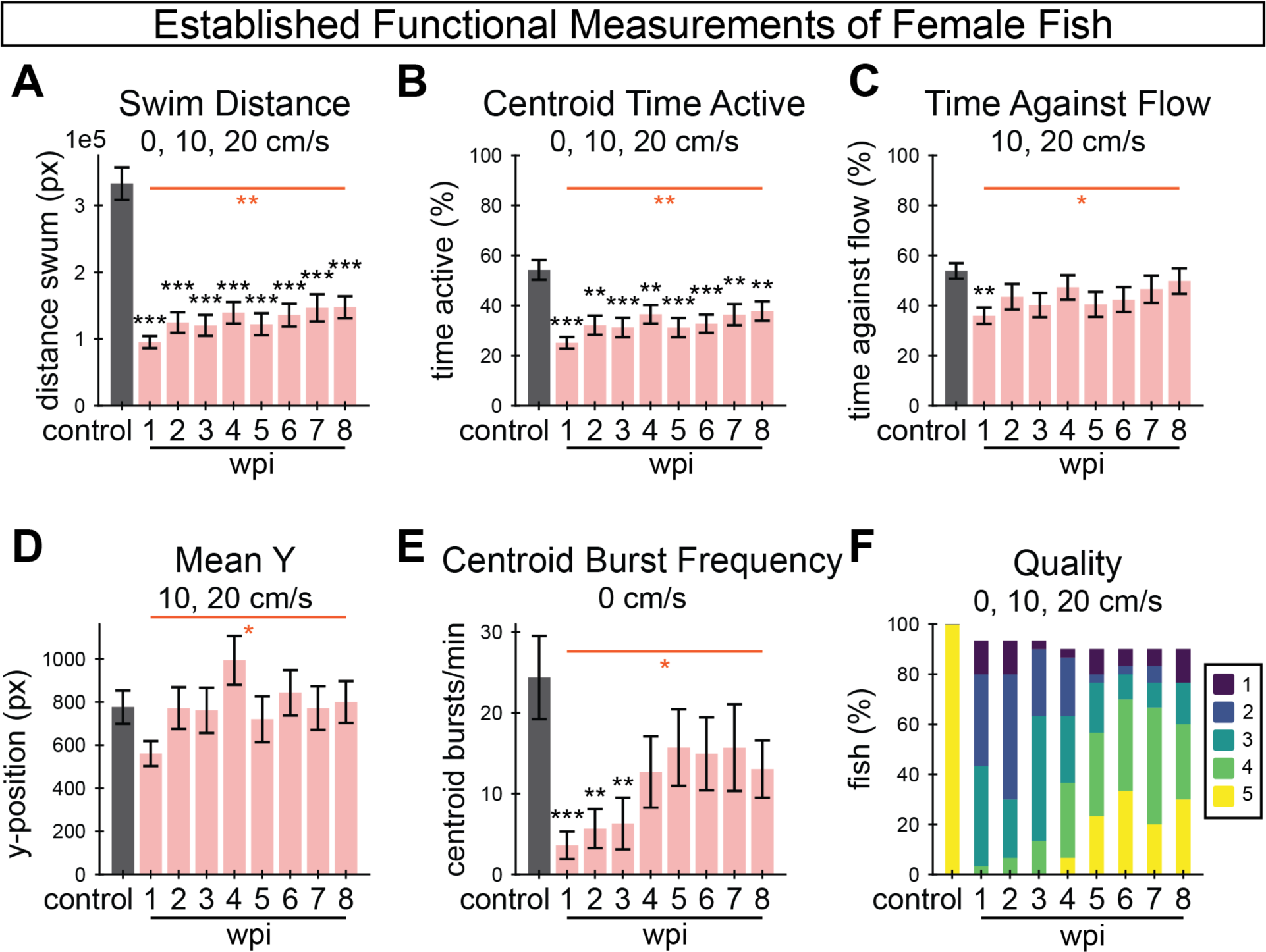
Established functional measurements for female fish in the tracking experiment. **(A-F)** A suite of established functional measurements were used to assess swim recovery for female fish following SCI. Swim distance (A), activity as measured by centroid movement (B), time swimming against the flow (C), mean y-position in the axis of flow (D), and burst frequency as measured by centroid movement (E) represent swim capacity measurements. Perceived swim quality scores are shown in F. Error bars depict SEM and statistical significance was determined by one-way repeated mixed measures ANOVA followed by post-hoc Student’s t-tests. p-value markers in black represent comparisons between each time point post-injury relative to control measurements prior to injury. Red horizontal bars and p-values marked in red show significance between 1 wpi and 8 wpi. ***P<0.001; **P<0.01; *P<0.05; ns, P>0.05.

**Figure S2.**
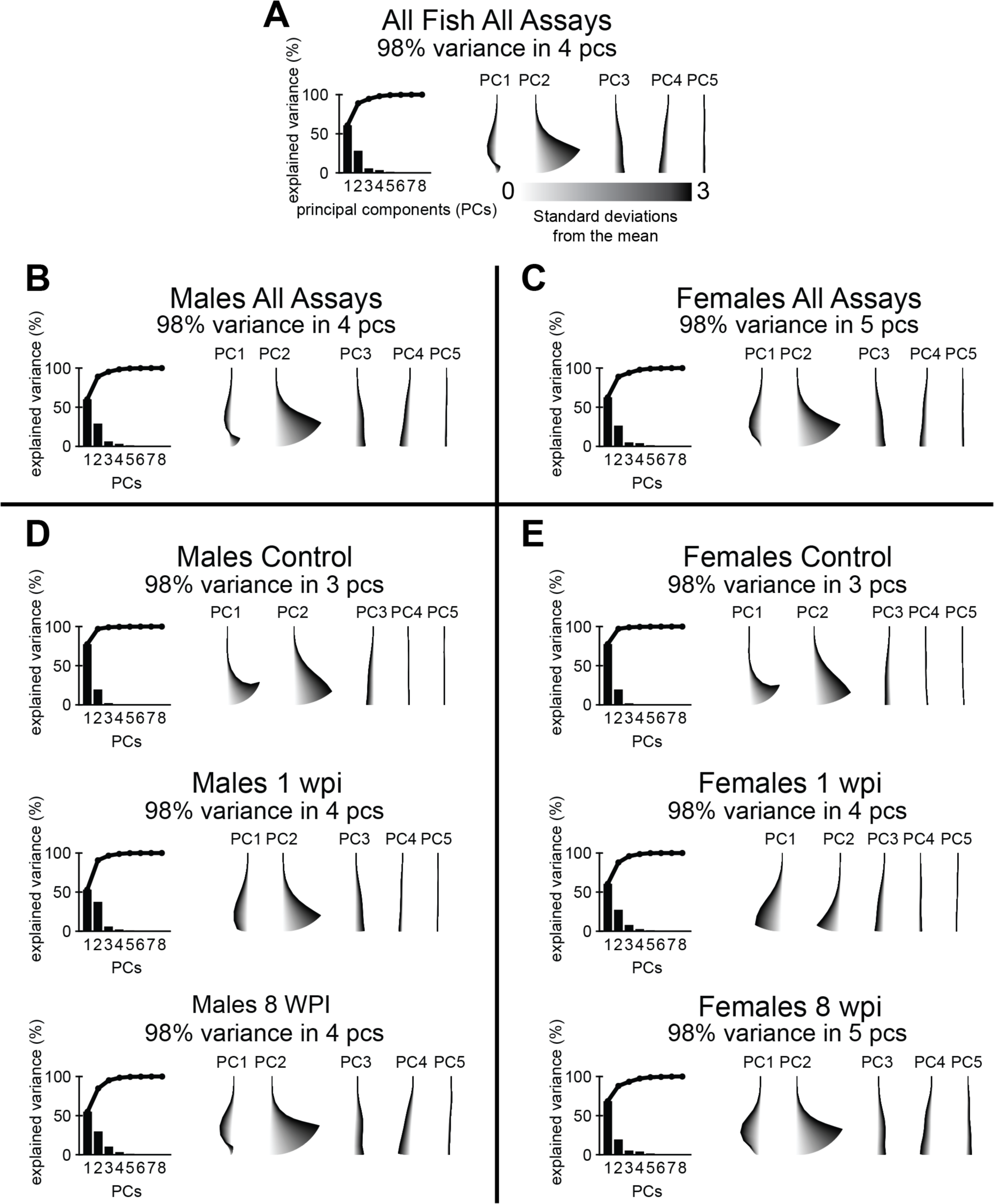
Principal component analysis of angle poses observed in the tracking experiment. On the left side of each PCA is a scree plot showing the percentage of variance captured by each PC. On the right side of each PCA are the first five components plotted as “eigenfish” by adding multiples of each PC to the mean pose. **(A)** A general representation of the posture space, and its complexity, for both healthy and injured fish. PCA of poses from all fish, all assays. **(B)** PCA of poses from all assays, male fish only. **(C)** PCA of poses from all assays, female fish only. **(D)** Posture space for male fish using poses taken from control, 1 wpi, and 8 wpi assays (top to bottom). **(E)** Posture space for female fish using poses taken from control, 1 wpi, and 8 wpi assays (top to bottom). Posture space is similarly complex whether separated by sex or taken altogether.

**Figure S3.**
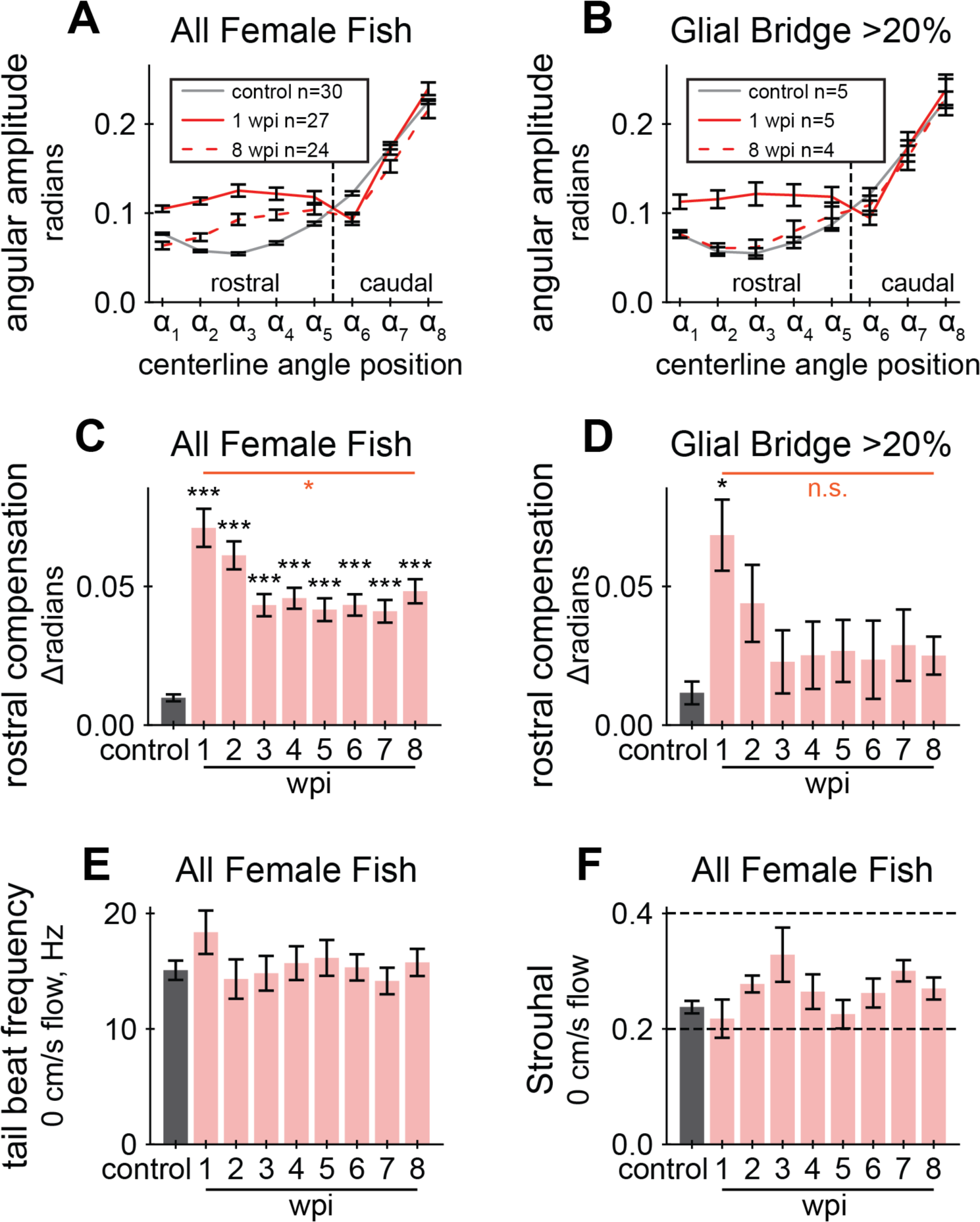
Cruise gait quality quantifications for female fish in the tracking experiment. **(A)** Cruise curvature profiles for female fish before injury, acutely injured fish at 1 wpi, and recovered fish at 8 wpi. Each cruise curvature profile represents the mean lateral angular amplitude along the dorsal centerline while cruising. The vertical dotted line separates rostral and caudal positions, demonstrating that acutely injured fish swim with markedly elevated curvature in the rostral portion of their body. **(B)** Cruise curvature profiles (control, 1wpi, and 8wpi) for female fish that regenerated more than 20% of the glial tissue at the lesion site. **(C)** Quantification of rostral compensation in female control fish prior to SC injury and at 1 to 8 wpi. Rostral compensation represents the displacement between cruise profiles. This score is the maximum distance, on the vertical axis, between rostral positions of a cruise profile from the control profile. Angle positions ⍺_1_ to ⍺_5_ from Panel D were used to define the rostral region. **(D)** Rostral compensation scores for female fish that regenerated more than 20% of the glial tissue at the lesion site. **(E)** Tail beat frequency for female fish in still water (0 cm/s), measured at each assayed week. ANOVA was not significant for tail beat frequencies. **(F)** Strouhal numbers for female fish in still water (0 cm/s), measured at each assayed week. Strouhal number is a unitless value related to vortex shedding mechanics and is defined as tail beat frequency times peak-to-peak amplitude of the tail tip divided by speed of forward motion. ANOVA was not significant for Strouhal numbers. Error bars depict SEM and statistical significance was determined by one-way repeated mixed measures ANOVA followed by post-hoc Student’s t-tests. p-value markers in black represent comparisons between each time point post-injury relative to control measurements prior to injury. Red horizontal bars and p-values marked in red show significance between 1 wpi and 8 wpi. ***P<0.001; **P<0.01; *P<0.05; ns, P>0.05.

**Figure S4.**
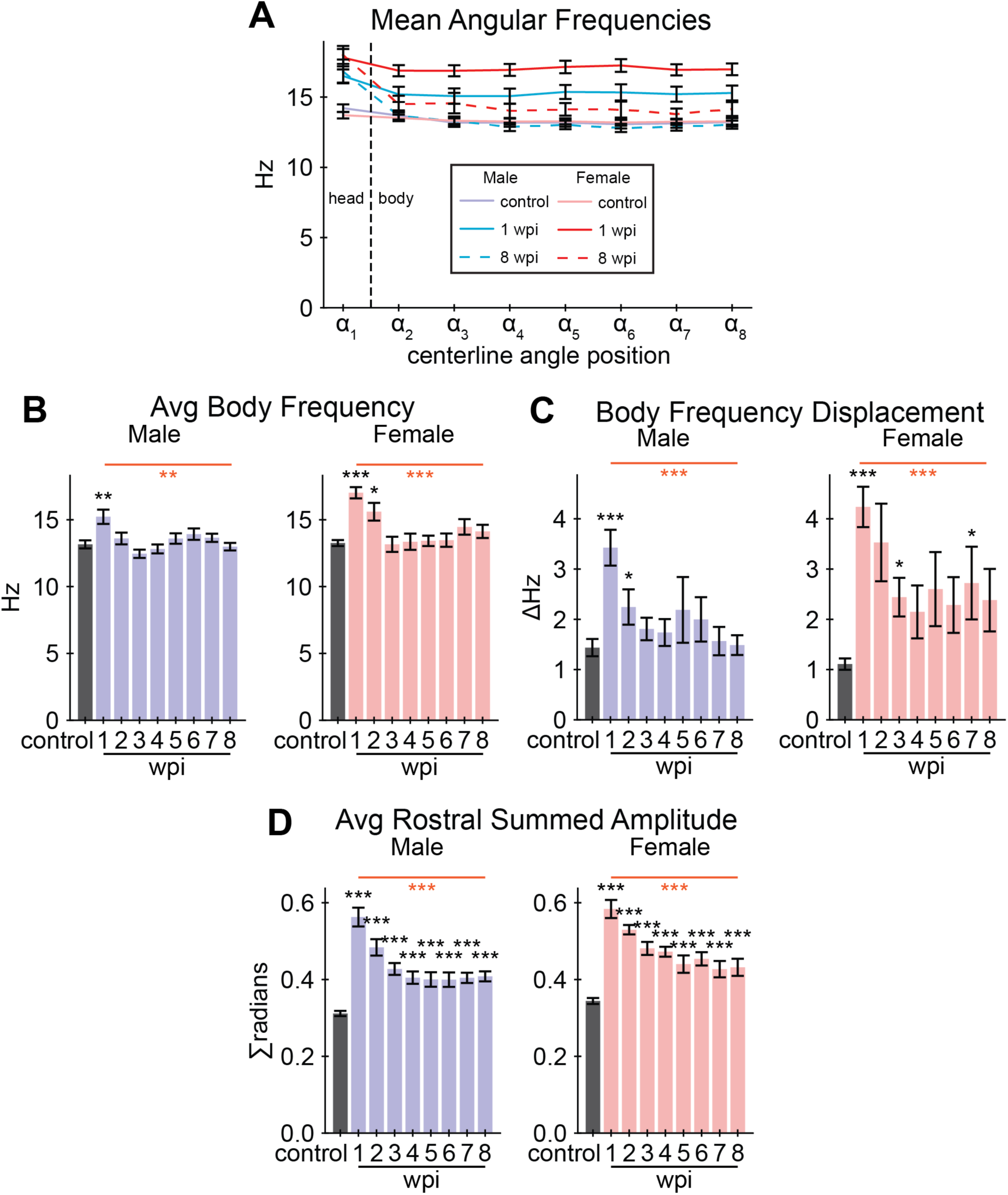
Cruise waveform features for male and female fish in the tracking experiment. We measured center-to-peak (half of peak-to-peak) curvature amplitude at all angle positions during cruise oscillations. We also measured cruise frequency as the number of complete oscillations at each angle position per second. The body of the fish corresponds to angle positions ⍺_2_ to ⍺_8_. The rostral region of the fish corresponds to angle positions ⍺_1_ to ⍺_5_. **(A)** Mean angle frequencies at each angle position on the centerline (x-axis) measured during cruise episodes, plotted for control, 1 wpi, and 8 wpi. Frequency was stable along the body of the centerline regardless of injury. **(B)** Average angular frequency along body positions for each assay and separated by sex. **(C)** Average body frequency displacement at each assay and separated by sex. Body frequency displacement is calculated similarly to rostral compensation: the maximum distance between a frequency curve and the control frequency curve at body positions. **(D)** The average sum of angular amplitudes measured at rostral positions along the fish, plotted for each assay and separated by sex. Error bars depict SEM and statistical significance was determined by one-way repeated mixed measures ANOVA followed by post-hoc Student’s t-tests. p-value markers in black represent comparisons between each time point post-injury relative to control measurements prior to injury. Red horizontal bars and p-values marked in red show significance between 1 wpi and 8 wpi. ***P<0.001; **P<0.01; *P<0.05; ns, P>0.05.

**Figure S5.**
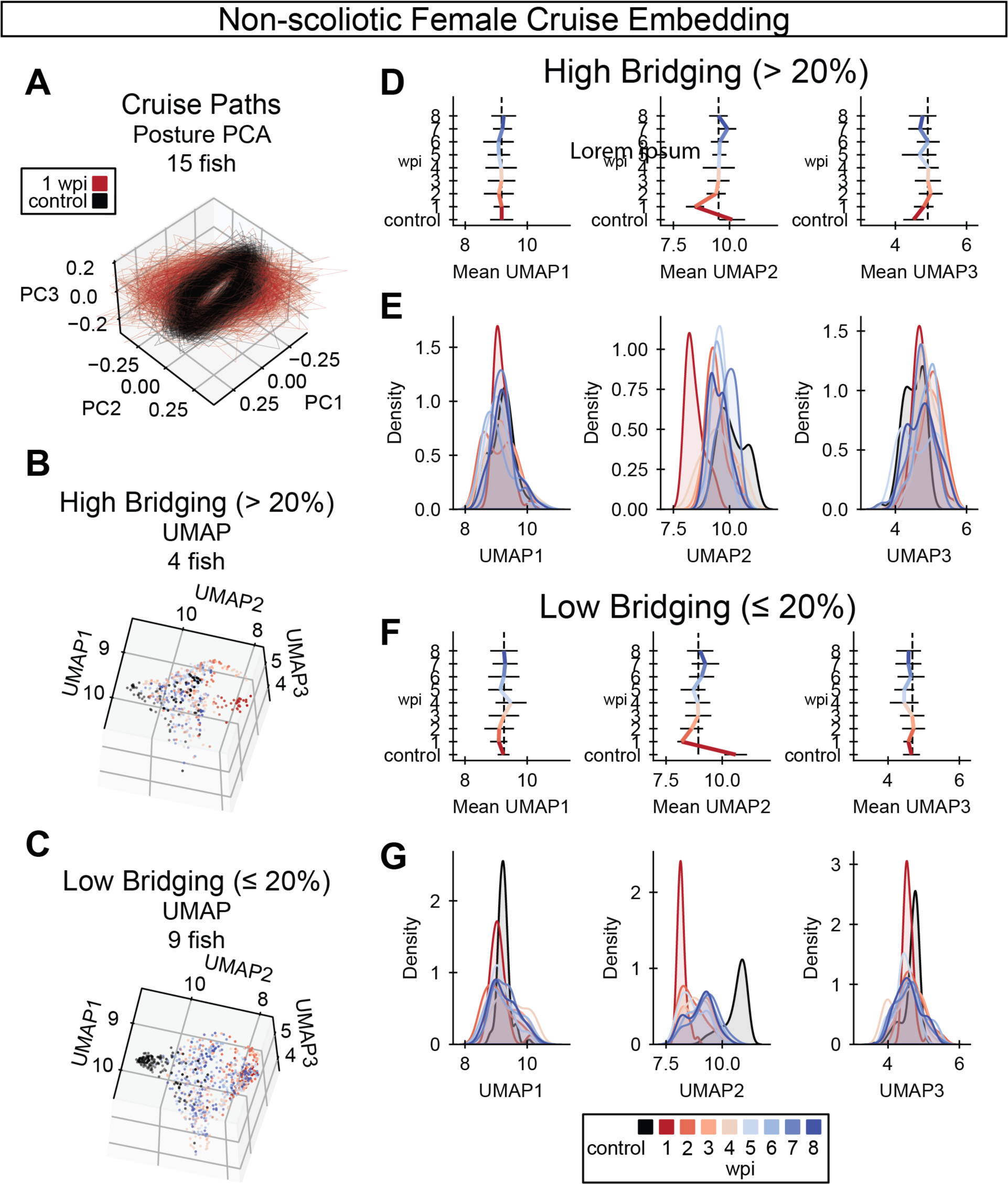
UMAP embedding of cruise behaviors during SC regeneration. **(A)** Cruise poses from control (black) and 1 wpi (red) female zebrafish were decomposed using principal component analysis (PCA). Female fish with scoliosis score < 0.35 at 8 wpi are shown. In this posture space, cruise trajectories from acutely injured fish at 1 wpi are markedly distinct from control fish trajectories. **(B-C)** Cruise poses were embedded using UMAP with precomputed dynamic time warping (DTW) distances. Cruise poses were analyzed for fish that regenerated well (glial bridging > 20%) in B and for fish with compromised cellular regeneration (glial bridging < 20%) in C. **(D)** Gait recovery trendlines in UMAP 1, 2, and 3 (left to right) for fish that regenerated well. Colored bars indicate the mean trend of each UMAP component. Error bars are standard deviations. The vertical dotted black lines align with the average UMAP value at 3 wpi. **(E)** Gaussian kernel density estimates for the regions of UMAP space occupied by cruises at each week post-injury for fish that regenerated well. The x-axes are matched between Panels D and E for comparison. **(F)** Gait recovery trendlines in UMAP 1, 2, and 3 (left to right) for fish with compromised cellular regeneration. Colored bars indicate the mean trend of each UMAP component. Error bars are standard deviations. The vertical dotted black lines align with the average UMAP value at 3 wpi. **(G)** Gaussian kernel density estimates for the regions of UMAP space occupied by cruises at each week post-injury for fish with compromised cellular regeneration. The x-axes are matched between Panels F and G for comparison.

**Figure S6.**
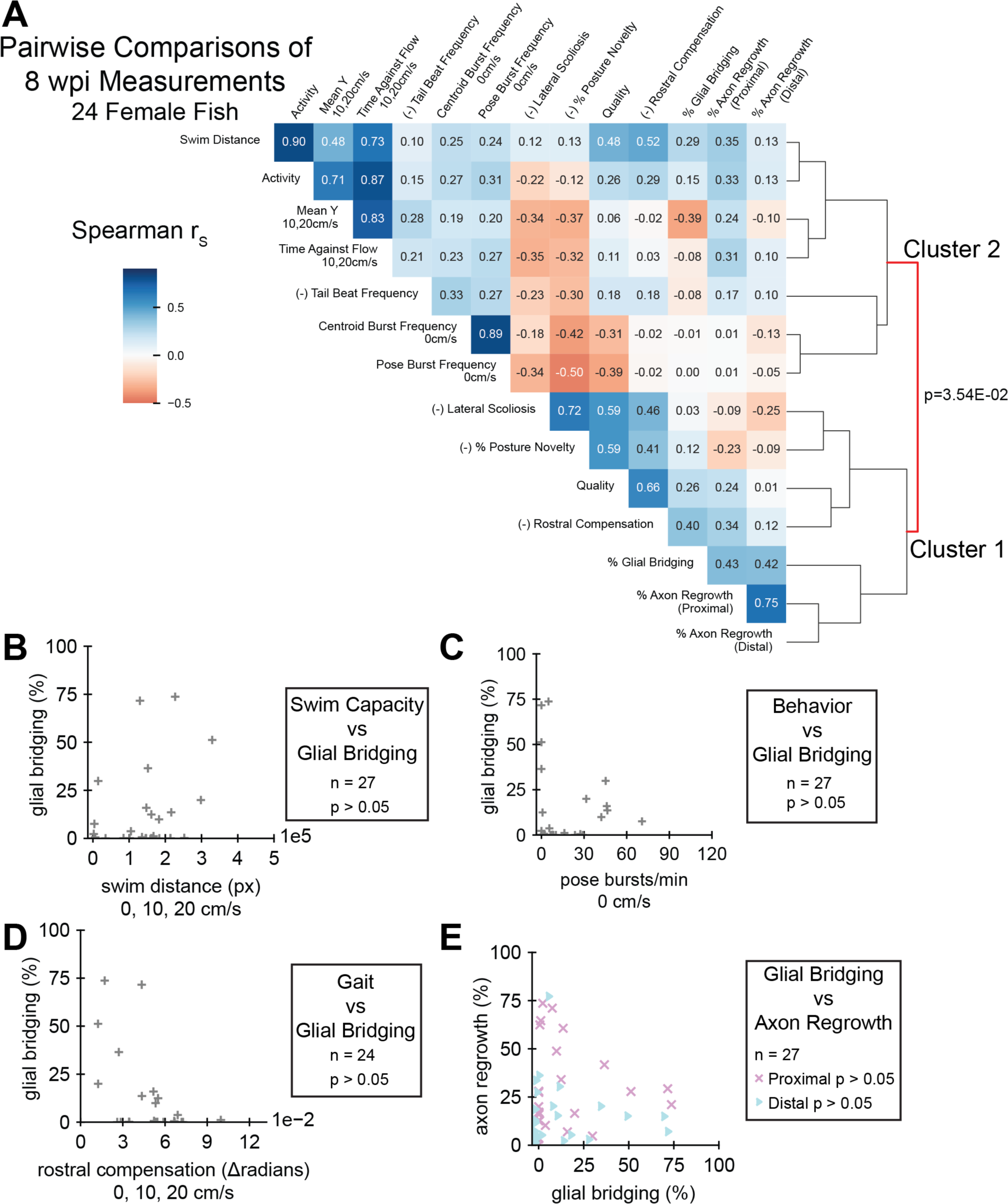
Pairwise comparisons of spinal cord regeneration metrics at 8 wpi. **(A)** Heatmap of Spearman’s rank correlations (r_s_) between the cellular and functional measurements taken. Regeneration metrics include swim capacity, swim quality, structural, and neurological measurements. A dendrogram representing similarities between correlation patterns of measurements is shown. Female fish at 8 wpi were analyzed. Fish that could not be measured in all attributes were omitted. Correlations are reported regardless of p-value. For the sake of meaningful clustering, a measured attribute was multiplied by -1 if its average was increased at 1 wpi compared to controls. These attributes are marked with the prefix “(-)” on the label. **(B-D)** One metric each of swim capacity (swim distance), behavior (burst frequency), and gait (rostral compensation) were plotted against glial bridging to demonstrate their relationship, though the correlation p-values were not significant. **(E)** Scatter plot showing a non-significant association between the size of regenerated glial tissue and axon regrowth, proximal and distal. The size of regenerated glial tissue plotted against axon regrowth, proximal and distal. r_s_ were not significant for Panels B-E, likely because most female fish in this experiment exhibited compromised glial regeneration.

**Figure S7.**
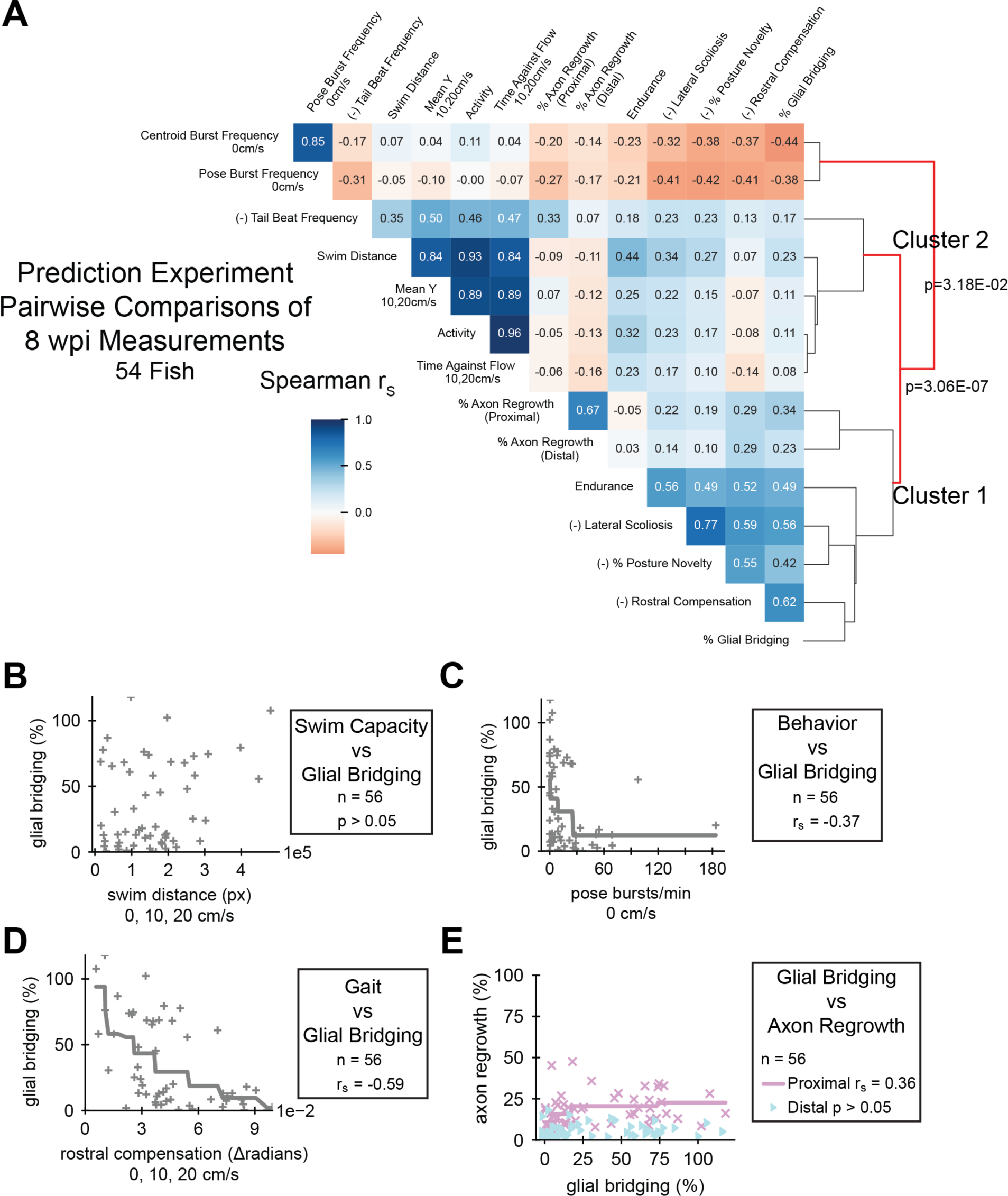
Prediction experiment: pairwise comparisons of spinal cord regeneration metrics at 8 wpi. **(A)** Heatmap of Spearman’s rank correlations (r_s_) between the cellular and functional measurements taken. Regeneration metrics include swim capacity, swim quality, structural, and neurological measurements. A dendrogram representing similarities between correlation patterns of measurements is shown. Male and female fish at 8 wpi were analyzed. Fish that could not be measured in all attributes were omitted. Correlations are reported regardless of p-value. For the sake of meaningful clustering, a measured attribute was multiplied by -1 if its average was increased at 1 wpi compared to controls. These attributes are marked with the prefix “(-)” on the label. **(B-D)** One metric each of swim capacity (swim distance), behavior (burst frequency), and gait (rostral compensation) were plotted against glial bridging to demonstrate their correlation. Because the Spearman correlation operates on ranked values, we fit monotonic splines to plots where r_s_ were significant (p < 0.05) to visualize possible associations. **(E)** Scatter plot showing a weak association between the size of regenerated glial tissue and axon regrowth, proximal and distal. Axon regrowth was measured lower than expected in this experiment.

## Material and Methods

### Zebrafish

Adult zebrafish of the Ekkwill and AB strains were maintained at the Washington University Zebrafish Core Facility. All animal experiments were performed in compliance with institutional animal protocols. Male and female animals between 3 and 9 months of ∼2–2.5 cm in length were used.

### SC transection

Zebrafish were anaesthetized using MS-222. Fine scissors were used to make a small incision that transects the spinal cord 4 mm caudal to the brainstem region. Complete transection was visually confirmed at the time of surgery. Injured animals were also assessed at 2 or 3 dpi to confirm loss of swim capacity post-surgery.

### Swim behavior recording

Two fish at a time were placed in a 5 L swim tunnel device (Loligo, cat# SW100605L, 120V/60Hz). A customized physical divider was secured in the center of the swim chamber parallel to the flow, separating the tunnel into 2 chambers to enable individual tracking of each fish. A camera was placed directly above the tunnel and recorded videos at 70 frames per second. Each 15-minute assay included three 5-minute periods of increasing water current velocity at 0, 10, and 20 cm/s.

### Swim endurance assay

The swim endurance assay performed in the prediction experiment was performed as previously described (40). Briefly, zebrafish were exercised in groups of 8–12 fish. After 10 minutes of acclimation inside the enclosed tunnel, water current velocity was increased every two minutes and fish swam against the current until they reached exhaustion. Exhausted animals were removed from the chamber without disturbing the remaining fish. Swim time and current velocity at exhaustion were recorded.

### Statistical analysis

Most statistical analysis was performed in Python (https://www.anaconda.com) version 3.7.3, with metric cluster significance (Figure 5, Figure S6, Figure S7) tested using R (46) version 4.0.4 and sigclust2 (47, 48) version 1.2.4. One-way repeated measures ANOVA and pairwise t-test post-hoc tests were performed using the package Pingouin (49) version 0.5.3. To test differences between outcome prediction groups, we used independent t-test in SciPy (50) version 1.7.3. Plots were generated using Matplotlib (51) version 3.4.3 and Seaborn (52) version 0.12.1, and were compiled and stylized using Adobe Illustrator.

### Software

All analysis was performed in Python (https://www.anaconda.com) version 3.7.3 and R (46) version 4.0.4. The primary packages used for our analyses include NumPy (53) version 1.21.5, pandas (54, 55) version 1.2.4, tensortools (56) version 0.3, scikit-learn (57) version 1.0.2, SciPy (50) version 1.7.3, Pingouin (49) version 0.5.3, dtw-python (58) version 1.1.12, umap-learn (59) version 0.4.2, DeepLabCut (24, 37) version 2.1.6.2, Matplotlib (51) version 3.4.3, Seaborn (52) version 0.12.1, sigclust2 (47, 48) version 1.2.4, and FFmpeg (https://ffmpeg.org) version 4.3.2.

### Video processing and pose annotation

We normalized each video and subtracted the background. Because a divider separated two fish in each assay, we separated the left and right sides of the video leaving one visible fish in each normalized video. We trained a DeepLabCut model to identify ten points on the dorsal centerline, from the head to the base of the tail fin, and a point near the tip of the tail fin (Figure 2C) (24, 37). To smooth the poses, we fit cubic splines to each computer-annotated pose and resampled ten evenly spaced points from the spline. We filtered away poses that were likely incorrectly annotated according to the following criteria: DeepLabCut likelihood score (require >0.5 at all keypoints), total pose length (require within 8 median absolute deviations from the median length), pose curvature (require z-score < 8 at each angle position), and whether the pose was too far from annotated poses in adjacent frames (require movement within 45 pixel at all keypoints).

Angles along the centerline were calculated using triplets of adjacent keypoints on the fish’s centerline (see keypoint and angle positions in Figure 2C). We called this sequence of angles an angle pose. For example, to calculate the angle at Point B from three sequential keypoints A, B, and C, the angle at B is how far Line AB must rotate to be parallel to Line BC. In this manner, each ten-keypoint pose was reduced to an angle pose with eight measured angles. Eight angle positions have been shown to be sufficient to capture ordinary variation in zebrafish posture (12). Although other research has used different methods to convert keypoints to angle poses, we chose this representation because the summed angle pose for a resting non-scoliotic fish is close to zero.

### Behavior annotation

We annotated many episodes of simplified behaviors: rest (1355 episodes), cruise (3243), and clockwise (128) and counterclockwise (165) turns. For the annotation model, we defined “cruise” as a forward swim with three body oscillations minimum. If the fish performed the behavior in the assay, we annotated three cruises per flow speed for every fish’s uninjured control, 1 week-post injury (wpi), 3 wpi, and 6 wpi assays. We annotated cruises in the same manner for all nine assays of 34 fish. Although we focused our analysis on cruise and rest behavior, we annotated and classified turns so the model would not mistakenly classify every non-rest behavior as cruise. Annotations included a nearly equal representation by sex (51% female). Representation by wpi was weighted to include more uninjured behaviors (pre: 26%, 1: 14%, 2: 6%, 3: 15%, 4: 4%, 5: 7%, 6: 14%, 7: 6%, 8: 8%).

To prepare the data for classification, episodes were split into frames. Each annotated frame’s angle pose was concatenated with the six angle poses before and the six angle poses after to create a larger vector, a temporal “pose window”, labeled with the middle frame’s annotated behavior. We withheld 25% of the annotated data for validation. Because cruises were overrepresented in our annotations, we oversampled the train and validation datasets such that all behaviors were equally represented with Synthetic Minority Over-sampling Technique (SMOTE) (60) using 20 nearest neighbors.

We fit a UMAP (59) model (2 components, 50 nearest neighbors) to all pose windows in the training set, then fit a random forest classifier (57) to the UMAP-transformed windows for final classification. Finally, because some assays contained unusually high numbers of rest frames incorrectly classified as active behaviors, particularly for severely scoliotic fish, we applied a posture change threshold: if an angle pose differed no more than a certain threshold from either the previous or the next pose, it was labeled as “rest”. We optimized this threshold using the same training set. Classifying and thresholding as described resulted in a high number of correctly classified cruises (94.8% training, 94.8% validation) and rests (96.5% training, 95.8% validation). We used this model to assign behavior labels to all frames with high-quality posture as previously described.

### Cruise embedding

Taking all annotated cruise episodes in each assay, we performed a 5-principal component decomposition on the angle poses, and then created a pairwise distance matrix for these cruise episodes using dynamic time warping. From each distance matrix, we used affinity propagation to select exemplars, or cruises that are most representative of the fish’s general behavior. If affinity propagation failed to converge, we randomly sampled at most 16 cruises from the assay. To calculate the embedding for several assays, we calculated a pairwise DTW distance matrix for all exemplar cruises from the assays, then fed the distance matrix into UMAP (59) using 20 nearest neighbors, which is consistent with published work (21). We omitted fish with scoliosis scores higher than 0.35.

### 8 wpi measurement correlation clustering

Measurements from 8 wpi assays were scaled to comparable ranges using StandardScaler from scikit-learn (57). Spearman correlations (r_s_) were calculated pairwise between measurements and a distance matrix of 1 – r_s_ was assembled using SciPy (50). Average linkage hierarchical clustering was performed, and significance between clusters was tested, using sigclust2, which was modified to accept 1 – r_s_ as the distance metric.

### TCA and fish clustering

A rank-3 tensor was created by concatenating all fish with their measurements in all assays (sorted temporally) in the tracking experiment. Measurements were scaled to comparable ranges using StandardScaler from scikit-learn (57). We omitted 6 fish that died before 8 wpi and 10 fish that did not perform a measurable cruise at every assayed week. Tensor component analysis was performed with tensortools (56). We compared ranks from 1 to 90 for three varieties of canonical polyadic (CP) tensor decompositions: unconstrained by alternating least squares (cp_als), nonnegative by block coordinate descent (ncp_bcd) and nonnegative by hierarchical alternating least squares (ncp_hals). After comparing model error and replicate similarity for 4 replicates of each rank and model, our chosen combination was rank-7 CP decomposition by alternating least squares (Figure 6A). We clustered fish according to their factor values using agglomerative clustering with distance threshold of 1.7 using scikit-learn. This distance threshold was chosen by comparing the number of clusters to the minimum cluster size for 1000 threshold values spaced evenly from 0.001 to 5 (Figure 6B).

### Measurements of functional and structural recovery

Classical Function

- Perceived Swim Quality
  - This score was assigned by observation on a scale from 1 (low quality) to 5 (high quality). We first found the best and worst fish, then scored all fish compared to these. We focused our observation on the first few minutes when the flow started (5 min) and when the flow accelerated (10 min). Our criteria were as follows: **5**: Fish’s swimming pattern looks normal. Mostly uses tail movement, not its head, to swim. No scoliosis. **4**: Fish seems like it can swim at the same rate and speed as those categorized as 5. Slight scoliosis. Slightly less tail movement than 5, and slightly more head movement or “wiggling”. **3**: Fish cannot swim as effectively as 4. Equal head and tail movement. Scoliosis looks “bent” at about a 45-degree angle. **2**: Fish have some tail movement, can rotate their body, but cannot swim forward effectively. Severe scoliosis with tail movement. No swimming once flow started. **1**: Fish have no tail movement. Severe scoliosis is observed. No swimming once flow started.
  - We treated the above criteria as guidelines because not all fish had all the features as described. For example, a fish could score “3” due to excessive head movement without significant scoliosis.
- Mean y-position
  - Average centroid position along the water flow axis in the swim tunnel. Higher “mean y” was closer to the flow source. It was assumed that exhausted fish would be pushed backward by the flow and have a lower mean-y. We reported mean y-position only for periods of the behavior assay when water was flowing (10, 20 cm/s).
- Distance
  - Sum of frame-to-frame centroid displacement measured in pixels. In this study we did not adjust distance according to water flow velocity, and we measured during the entire assay at all flow speeds.
- Activity
  - Sum of frames where centroid displacement (see Distance) was above a threshold, divided by the total number of frames. In this study we did not adjust activity according to water flow velocity, and we measured during the entire assay at all flow speeds.
- Centroid Burst Frequency
  - We defined a centroid burst as an uninterrupted period of active frames (see Activity). Centroid burst frequency is the number of centroid bursts per minute. We reported this measurement only for periods of the behavior assay when water was not flowing (0 cm/s).
- Time Against Flow
  - Sum of frames where centroid displacement (see Distance) is above a threshold in the direction of the flow, allowing 45-degree deviation in either direction, divided by the total number of frames where water is flowing. Accordingly, time against flow was only calculated for periods of the behavior assay when water was flowing (10, 20 cm/s).
- Endurance or Time to Exhaustion
  - The duration of time that a fish can swim against flowing water until exhausted. Fish allow the fish to acclimate in calm water, then increase the flow speed periodically. When the fish can no longer pull away from a barrier at the back of the tunnel, it is exhausted.

Gait and Posture

- Pose Burst Frequency
  - Same as Centroid Burst Frequency but active frames were defined as those with active posture; that is, for frames not classified as “rest” by our behavior classifier. Although this calculation for activity did not depend on flow velocity, we only reported this value for periods without water flow (0 cm/s) to be comparable with our reported centroid burst frequency.
- Lateral Scoliosis
  - For each period of rest, or inactive posture, we calculated the average sum of each angle pose, then took the weighted average of these values over the entire swim where weights are the length of the rest periods. Although the calculation would have been essentially equivalent for non-scoliotic fish, we chose to analyze each rest bout separately and average their results rather than taking the average posture over the entire swim because some fish with severely poor recovery could not control their dorsal-ventral orientation. For example, when a scoliotic fish flipped to its back, its posture inverted; and repeated flipping lead to an artificially lower scoliosis score when posture was averaged over the entire swim. Hence, each period of rest was analyzed separately, then values for all rest periods were averaged as described.
- Cruise Angular Frequency
  - The number of posture oscillations per second while swimming forward. Angular frequency was mostly uniform along the centerline (Figure S4A). We computed cruise angular frequency for each of the eight computed angle positions along a fish’s centerline, then averaged over the cruise. All cruise values were then averaged to get a single score for each assay. We reported average cruise angular frequency values in Figure S4B and displacement from control average in Figure S4C.
- Cruise Rostral Compensation
  - First, we calculated average angular amplitudes at each angle position along the dorsal centerline during episodes of cruise behavior. We called this a cruise curvature profile. We calculated a control profile as the average profile of uninjured fish. We defined rostral compensation as the maximum absolute difference between an assay’s cruise curvature profile and the control profile at rostral angle positions ⍺_1_ to ⍺_5_ (Figure 2C).
- Posture Novelty
  - We trained a Local Outlier Factor (LOF) model using sampled uninjured control poses, then used the model to perform novelty detection on all assays. An assay’s posture novelty score was calculated as the number of poses identified as “novel” by LOF divided by the total number of poses in the assay.
  - We created a representative training set by sampling 1200 poses from each control assay: 100 poses from 12 k-mean clusters. K-means clustering was performed separately for each assay.
- Tail Beat Frequency
  - Using an annotated point near the tip of the caudal fin, we measured lateral extrema, then calculated oscillation frequency using those measurements.
- Strouhal Number
  - We calculate tail beat frequencies and peak-to-peak caudal fin tip displacement amplitudes for each cruise episode. Strouhal number (St) was defined as St = ƒ*A/U*, which is tail beat frequency (ƒ) times amplitude (*A*) divided by forward speed (*U*) (38). To get average St for one assay, we calculated St for all cruise episodes then took the average of these numbers weighted by the number of periods in each respective cruise.

Spinal Cord Structure

- % Bridging
  - GFAP immunohistochemistry was performed on serial transverse sections as previously described(40). The cross-sectional area of the glial bridge and the area of the intact SC rostral to the lesion were measured using ImageJ (Fiji) software(61). Bridging was calculated as a ratio of these measurements.
- % Axon Regrowth Proximal and Distal
  - Anterograde axon tracing was performed as previously described(40). Fish were anaesthetized using MS-222 and fine scissors were used to transect the cord 4 mm rostral to the lesion site. Biocytin-soaked Gelfoam Gelatin Sponge was applied at the new injury site (Gelfoam, Pfizer, cat# 09-0315-08; Biocytin, saturated solution, Sigma, cat# B4261). Fish were euthanized 6 hours post-treatment and Biocytin-labeled axons were histologically detected using Alexa Fluor 594-conjugated Streptavidin (Thermo Fisher, cat# S-11227). Biocytin-labeled axons were quantified using the “threshold” and “particle analysis” tools in the Fiji software. Four sections per fish at 0.5 (proximal) and 2 (distal) mm caudal to the lesion core, and 2 sections 1 mm rostral to the lesion, were analyzed. Regrowth was calculated as a ratio of these measurements.

## Bibliography

1. Cigliola V, Becker CJ, Poss KD. Building bridges, not walls: spinal cord regeneration in zebrafish. Disease Models & Mechanisms. 2020;13(5):dmm044131.

2. Mokalled MH, Patra C, Dickson AL, Endo T, Stainier DYR, Poss KD. Injury-induced ctgfa directs glial bridging and spinal cord regeneration in zebrafish. Science. 2016;354(6312):630–4.

3. Becker T, Wullimann MF, Becker CG, Bernhardt RR, Schachner M. Axonal regrowth after spinal cord transection in adult zebrafish. The Journal of Comparative Neurology. 1997;377(4):577–95.

4. Goldshmit Y, Sztal TE, Jusuf PR, Hall TE, Nguyen-Chi M, Currie PD. Fgf-dependent glial cell bridges facilitate spinal cord regeneration in zebrafish. The Journal of Neuroscience. 2012;32(22):7477–92.

5. Strand NS, Hoi KK, Phan TMT, Ray CA, Berndt JD, Moon RT. Wnt/β-catenin signaling promotes regeneration after adult zebrafish spinal cord injury. Biochemical and biophysical research communications. 2016;477(4):952–6.

6. van Raamsdonk W, Maslam S, de Jong DH, Smit-Onel MJ, Velzing E. Long term effects of spinal cord transection in Zebrafish: swimming performances, and metabolic properties of the neuromuscular system. Acta Histochemica. 1998;100(2):117–31.

7. Becker CG, Lieberoth BC, Morellini F, Feldner J, Becker T, Schachner M. L1.1 is involved in spinal cord regeneration in adult zebrafish. The Journal of Neuroscience. 2004;24(36):7837–42.

8. Klatt Shaw D, Mokalled MH. Efficient CRISPR/Cas9 mutagenesis for neurobehavioral screening in adult zebrafish. G3 (Bethesda, Md). 2021;11(8).

9. Burris B, Jensen N, Mokalled MH. Assessment of swim endurance and swim behavior in adult zebrafish. Journal of Visualized Experiments. 2021(177).

10. Saraswathy VM, Zhou L, Mcadow AR, Burris B, Dogra D, Reischauer S, et al. Myostatin is a negative regulator of adult neurogenesis after spinal cord injury in zebrafish. Cell Reports. 2022;41(8):111705.

11. Huang C-X, Zhao Y, Mao J, Wang Z, Xu L, Cheng J, et al. An injury-induced serotonergic neuron subpopulation contributes to axon regrowth and function restoration after spinal cord injury in zebrafish. Nature Communications. 2021;12(1):7093.

12. Huang C-X, Wang Z, Cheng J, Zhu Z, Guan NN, Song J. De novo establishment of circuit modules restores locomotion after spinal cord injury in adult zebrafish. Cell Reports. 2022;41(4):111535.

13. Chang W, Pedroni A, Bertuzzi M, Kizil C, Simon A, Ampatzis K. Locomotion dependent neuron-glia interactions control neurogenesis and regeneration in the adult zebrafish spinal cord. Nature Communications. 2021;12(1).

14. Hosseini P, Mirsadeghi S, Rahmani S, Izadi A, Rezaei M, Ghodsi Z, et al. Dopamine Receptors Gene Expression Pattern and Locomotor Improvement Differ Between Female and Male Zebrafish During Spinal Cord Auto Repair. Zebrafish. 2022;19(4):137–47.

15. Zeng C-W, Kamei Y, Shigenobu S, Sheu J-C, Tsai H-J. Injury-induced Cavl-expressing cells at lesion rostral side play major roles in spinal cord regeneration. Open biology. 2021;11(2):200304.

16. Vasudevan D, Liu Y-C, Barrios JP, Wheeler MK, Douglass AD, Dorsky RI. Regenerated interneurons integrate into locomotor circuitry following spinal cord injury. Experimental Neurology. 2021;342:113737.

17. Hossainian D, Shao E, Jiao B, Ilin VA, Parris RS, Zhou Y, et al. Quantification of functional recovery in a larval zebrafish model of spinal cord injury. Journal of Neuroscience Research. 2022.

18. Doyle LMF, Roberts BL. Exercise enhances axonal growth and functional recovery in the regenerating spinal cord. Neuroscience. 2006;141(1):321–7.

19. Ljunggren EE, Haupt S, Ausborn J, Ampatzis K, El Manira A. Optogenetic Activation of Excitatory Premotor Interneurons Is Sufficient to Generate Coordinated Locomotor Activity in Larval Zebrafish. Journal of Neuroscience. 2014;34(1):134–9.

20. Girdhar K, Gruebele M, Chemla YR. The behavioral space of zebrafish locomotion and its neural network analog. Plos One. 2015;10(7):e0128668.

21. Mearns DS, Donovan JC, Fernandes AM, Semmelhack JL, Baier H. Deconstructing Hunting Behavior Reveals a Tightly Coupled Stimulus-Response Loop. Current Biology. 2020;30(1):54–69.e9.

22. Fontaine E, Lentink D, Kranenbarg S, Müller UK, van Leeuwen JL, Barr AH, et al. Automated visual tracking for studying the ontogeny of zebrafish swimming. Journal of Experimental Biology. 2008;211(Pt 8):1305–16.

23. Thomas LSV, Gehrig J. Multi-template matching: a versatile tool for object-localization in microscopy images. BMC Bioinformatics. 2020;21(1).

24. Mathis A, Mamidanna P, Cury KM, Abe T, Murthy VN, Mathis MW, et al. DeepLabCut: markerless pose estimation of user-defined body parts with deep learning. Nature Neuroscience. 2018;21(9):1281–9.

25. Graving JM, Chae D, Naik H, Li L, Koger B, Costelloe BR, et al. DeepPoseKit, a software toolkit for fast and robust animal pose estimation using deep learning. eLife. 2019;8.

26. Liu X, Yu S-Y, Flierman NA, Loyola S, Kamermans M, Hoogland TM, et al. OptiFlex: Multi-Frame Animal Pose Estimation Combining Deep Learning With Optical Flow. Frontiers in Cellular Neuroscience. 2021;15:621252.

27. Pereira TD, Tabris N, Matsliah A, Turner DM, Li J, Ravindranath S, et al. SLEAP: A deep learning system for multi-animal pose tracking. Nature Methods. 2022;19(4):486–95.

28. Mwaffo V, Zhang P, Romero Cruz S, Porfiri M. Zebrafish swimming in the flow: a particle image velocimetry study. PeerJ. 2017;5:e4041.

29. McCullough MH, Goodhill GJ. Unsupervised quantification of naturalistic animal behaviors for gaining insight into the brain. Current Opinion in Neurobiology. 2021;70:89–100.

30. Hsu AI, Yttri EA. B-SOiD, an open-source unsupervised algorithm for identification and fast prediction of behaviors. Nature Communications. 2021;12(1).

31. Sun G, Lyu C, Cai R, Yu C, Sun H, Schriver KE, et al. DeepBhvTracking: A novel behavior tracking method for laboratory animals based on deep learning. Frontiers in Behavioral Neuroscience. 2021;15:750894.

32. Bohnslav JP, Wimalasena NK, Clausing KJ, Dai YY, Yarmolinsky DA, Cruz T, et al. DeepEthogram, a machine learning pipeline for supervised behavior classification from raw pixels. eLife. 2021;10.

33. Xu J, Hu C, Jiang Q, Pan H, Shen H, Schachner M. Trimebutine, a small molecule mimetic agonist of adhesion molecule L1, contributes to functional recovery after spinal cord injury in mice. Disease Models & Mechanisms. 2017;10(9):1117–28.

34. Vrinten DH, Hamers FFT. ’CatWalk’ automated quantitative gait analysis as a novel method to assess mechanical allodynia in the rat; a comparison with von Frey testing. Pain. 2003;102(1-2):203–9.

35. Barbeau H, Ladouceur M, Norman KE, Pépin A, Leroux A. Walking after spinal cord injury: evaluation, treatment, and functional recovery. Archives of Physical Medicine and Rehabilitation. 1999;80(2):225–35.

36. Maynard FM, Bracken MB, Creasey G, Donovan WH, Ducker TB, Garber SL, et al. International standards for neurological and functional classification of spinal cord injury. Spinal cord. 1997;35(5):266–74.

37. Nath T, Mathis A, Chen AC, Patel A, Bethge M, Mathis MW. Using DeepLabCut for 3D markerless pose estimation across species and behaviors. Nature Protocols. 2019;14(7):2152–76.

38. Taylor GK, Nudds RL, Thomas ALR. Flying and swimming animals cruise at a Strouhal number tuned for high power efficiency. Nature. 2003;425(6959):707–11.

39. Delcourt J, Ovidio M, Denoël M, Muller M, Pendeville H, Deneubourg J-L, et al. Individual identification and marking techniques for zebrafish. Reviews in Fish Biology and Fisheries. 2018;28(4):839–64.

40. Klatt Shaw D, Saraswathy VM, Zhou L, Mcadow AR, Burris B, Butka E, et al. Localized EMT reprograms glial progenitors to promote spinal cord repair. Developmental Cell. 2021;56(5):613–26.e7.

41. Beal DN, Hover FS, Triantafyllou MS, Liao JC, Lauder GV. Passive propulsion in vortex wakes. Journal of fluid mechanics. 2006;549(-1):385.

42. Barrios JP, Wang W-C, England R, Reifenberg E, Douglass AD. Hypothalamic dopamine neurons control sensorimotor behavior by modulating brainstem premotor nuclei in zebrafish. Current Biology. 2020;30(23):4606–18.e4.

43. Bagnall MW, McLean DL. Modular organization of axial microcircuits in zebrafish. Science. 2014;343(6167):197–200.

44. Reimer MM, Sörensen I, Kuscha V, Frank RE, Liu C, Becker CG, et al. Motor neuron regeneration in adult zebrafish. The Journal of Neuroscience. 2008;28(34):8510–6.

45. Kathe C, Skinnider MA, Hutson TH, Regazzi N, Gautier M, Demesmaeker R, et al. The neurons that restore walking after paralysis. Nature. 2022;611(7936):540–7.

46. Team RC. R: A Language and Environment for Statistical Computing. Vienna, Austria 2021.

47. Kimes PK, Liu Y, Neil Hayes D, Marron JS. Statistical significance for hierarchical clustering. Biometrics. 2017;73(3):811–21.

48. Kimes P. sigclust2: sigclust2: Statistical Significance of Clustering. 2018.

49. Vallat R. Pingouin: statistics in Python. Journal of Open Source Software. 2018;3(31):1026.

50. Virtanen P, Gommers R, Oliphant TE, Haberland M, Reddy T, Cournapeau D, et al. SciPy 1.0: fundamental algorithms for scientific computing in Python. Nature Methods. 2020;17(3):261–72.

51. Hunter JD. Matplotlib: A 2D Graphics Environment. Computing in Science & Engineering. 2007;9(3):90–5.

52. Waskom M. seaborn: statistical data visualization. Journal of Open Source Software. 2021;6(60):3021.

53. Harris CR, Millman KJ, Van Der Walt SJ, Gommers R, Virtanen P, Cournapeau D, et al. Array programming with NumPy. Nature. 2020;585(7825):357–62.

54. The pandas development t. pandas-dev/pandas: Pandas 1.0.3. Zenodo. 2020.

55. McKinney W, editor Data structures for statistical computing in python. Python in Science Conference; 2010 2010: SciPy.

56. Williams AH, Kim TH, Wang F, Vyas S, Ryu SI, Shenoy KV, et al. Unsupervised Discovery of Demixed, Low-Dimensional Neural Dynamics across Multiple Timescales through Tensor Component Analysis. Neuron. 2018;98(6):1099–115.e8.

57. Pedregosa F, Varoquaux G, Gramfort A, Michel V, Thirion B, Grisel O, et al. Scikit-learn: Machine Learning in Python. Journal of Machine Learning Research. 2011;12:2825–30.

58. Giorgino T. Computing and Visualizing Dynamic Time Warping Alignments in R: The dtw Package. Journal of Statistical Software. 2009;31(7):1–24.

59. McInnes L, Healy J, Saul N, Großberger L. UMAP: uniform manifold approximation and projection. The Journal of Open Source Software. 2018;3(29):861.

60. Lemaitre G, Nogueira F, Aridas CK. Imbalanced-learn: A Python Toolbox to Tackle the Curse of Imbalanced Datasets in Machine Learning. Journal of Machine Learning Research. 2017;18:1–5.

61. Schindelin J, Arganda-Carreras I, Frise E, Kaynig V, Longair M, Pietzsch T, et al. Fiji: an open-source platform for biological-image analysis. Nature Methods. 2012;9(7):676–82.

